# Mapping DNA sequence to transcription factor binding energy *in vivo*

**DOI:** 10.1101/331124

**Authors:** Stephanie L. Barnes, Nathan M. Belliveau, William T. Ireland, Justin B. Kinney, Rob Phillips

## Abstract

Despite the central importance of transcriptional regulation in systems biology, it has proven difficult to determine the regulatory mechanisms of individual genes, let alone entire gene networks. It is particularly difficult to analyze a promoter sequence and identify the locations, regulatory roles, and energetic properties of binding sites for transcription factors and RNA polymerase. In this work, we present a strategy for interpreting transcriptional regulatory sequences using *in vivo* methods (i.e. the massively parallel reporter assay Sort-Seq) to formulate quantitative models that map a transcription factor binding site’s DNA sequence to transcription factor-DNA binding energy. We use these models to predict the binding energies of transcription factor binding sites to within 1 *k*_*B*_*T* of their measured values. We further explore how such a sequence-energy mapping relates to the mechanisms of trancriptional regulation in various promoter contexts. Specifically, we show that our models can be used to design specific induction responses, analyze the effects of amino acid mutations on DNA sequence preference, and determine how regulatory context affects a transcription factor’s sequence specificity.

## Introduction

High-throughput sequencing allows us to sequence the genome of nearly any species at will. The amount of genomic data available is already enormous and will only continue to grow. However, this mass of data is largely uninformative without appropriate methods of analyzing it. Despite decades of research, much genomic data still defies our efforts to interpret it. It is particularly challenging to interpret non-coding DNA such as intergenic regulatory regions. We can infer the locations of some transcription start sites and transcription factor binding sites, but these inferences tell us little about the functional role of these putative sites. In order to better interpret these types of sequences, we need a better understanding of how sequence elements control gene expression. An important avenue for developing this level of understanding is to propose models that map sequence to function and to perform experiments that test these models.

Over half of the genes in *E. coli*, which is arguably the best-understood model organism, lack any regulatory annotation (see RegulonDB [1]). Those operons whose regulation is well described (e.g. the *lac, rel*, and *mar* operons [2–4]) required decades of study involving laborious genetic and biochemical experiments [5]. A wide variety of new techniques have been proposed and implemented to simplify the process of determining how a gene is regulated. Chromatin immunoprecipitation (ChIP) based methods such as ChIP-chip and ChIP-seq make it possible to determine the genome-wide binding locations of individual transcription factors of interest. Massively parallel reporter assays (MPRAs) have made it possible to read out transcription factor binding position and occupancy *in vivo* with base-pair resolution, and provide a means for analyzing additional features such as “insulator” sequences [6–8]. *In vitro* methods based on protein-binding microarrays [9], SELEX [10–12], MITOMI [13–15], and binding assays performed in high-throughput sequencing flow cells [16,17] have made it possible to measure transcription factor affinity to a broad array of possible binding sites and can also account for features such as flanking sequences [15, 18, 19]. However, *in vitro* methods cannot fully account for the *in vivo* consequences of binding site context and interactions with other proteins. Current *in vivo* methods for measuring transcription factor binding affinities, such as bacterial one-hybrid [20, 21], require a restructuring of the promoter so that it no longer resembles its genomic counterpart. Additionally, efforts to computationally ascertain the locations of transcription factor binding sites frequently produce false positives [22, 23]. Furthermore, a common assumption underlying many of these methods is that transcription factor occupancy in the vicinity of a promoter implies regulation, but it has been shown that occupancy cannot always accurately predict the effect of a transcription factor on gene regulation [24,25]. As these examples show, it remains challenging to integrate multiple aspects of transcription factor binding into a cohesive understanding of gene regulation.

Here we work to develop such a cohesive understanding by integrating rigorous thermodynamic modeling with *in vivo* transcription factor binding experiments. In Ref. [26], we showed that the MPRA Sort-Seq [27], combined with a simple linear model for protein-DNA binding specificity, can be used to accurately predict the binding energies of multiple RNAP binding site mutants, serving as a jumping off point for the use of such models as a quantitative tool in synthetic biology. Here we apply this technique to transcription factor binding sites in an effort to better understand how transcription factors interact with regulatory DNA under different conditions. Specifically, we use Sort-Seq to map sequence to binding energy for a repressor-operator interaction, and we rigorously characterize the variables that must be considered in order to obtain an accurate mapping between DNA sequence and binding energy. We then use our sequence-energy mapping to design a series of operators with a hierarchy of controlled binding energies measured in *k*_*B*_*T* units. To demonstrate our control over these operators and their associated regulatory logic, we use these characterized binding sites to design a wide range of induction responses with different phenotypic properties such as leakiness, dynamic range and [*EC*_50_]. Next, we focus our attention on the synergy between mutations in the amino acid sequence of transcription factors and their corresponding binding sites. Finally, we show the broader reach of these results by exploring how binding site position and regulatory context can change the DNA-protein sequence specificity for multiple different transcription factors.

## Results

### Obtaining energy matrices using Sort-Seq

A major goal of this study was to show that one can use Sort-Seq to precisely map DNA sequence to binding energy for a transcription factor binding site, thus making it possible to predict and manipulate transcriptional activity *in vivo*. While numerous *in vitro* studies have successfully mapped sequence to affinity [9–17], and some *in vivo* studies have used methods such as bacterial one-hybrid to provide such mappings as well [20, 21], these studies are limited because they do not reflect the actual wild-type arrangement of regulatory elements, thus potentially missing vital regulatory information. Moreover, while position-weight matrices (PWMs) derived from genomic data have traditionally been used to ascertain *in vivo* sequence specificities, it can be difficult to convert these specificities into quantitative binding energy mappings due to the relatively small number of sequences that are used to generate these PWMs.

Sort-Seq has previously been shown to be a promising technique for mapping protein binding sequences to binding energies. In Ref. [26], binding energy predictions for RNAP were made from an energy matrix generated in Ref. [27] that used the wild-type *lac* promoter as a reference sequence (i.e. the sequence that was mutated to perform Sort-Seq). Here, we design experiments that use the Sort-Seq technique described in [27] with the specific intent of creating energy matrices with maximum predictive power (see Fig 1), and we test the predictions from these matrices against measured binding energies. We show that such predictive matrices can be produced for multiple transcription factors (e.g. XylR, PurR, and LacI) implicated in an array of regulatory architectures. To thoroughly test the accuracy of our predictive matrices, we begin with promoters that employ “simple repression,” in which a repressor binds to an operator such that it occludes RNAP binding, thereby preventing transcription and repressing the gene [28]. As a model for how sequence-energy mappings might be used for transcription factor binding sites in simple repression architectures, we interrogate the binding specificity of the *lac* repressor (LacI). LacI was chosen for this role because it is well-characterized and has known binding sites in only one operon within the genome, making it an ideal choice for this kind of systematic and rigorous analysis. We create three distinct energy matrices in which each of the natural *lac* operators (O1, O2, or O3 [2]) acts as the reference sequence. Appendix A lists the wild-type sequences for these simple repression constructs.

**Figure 1.**
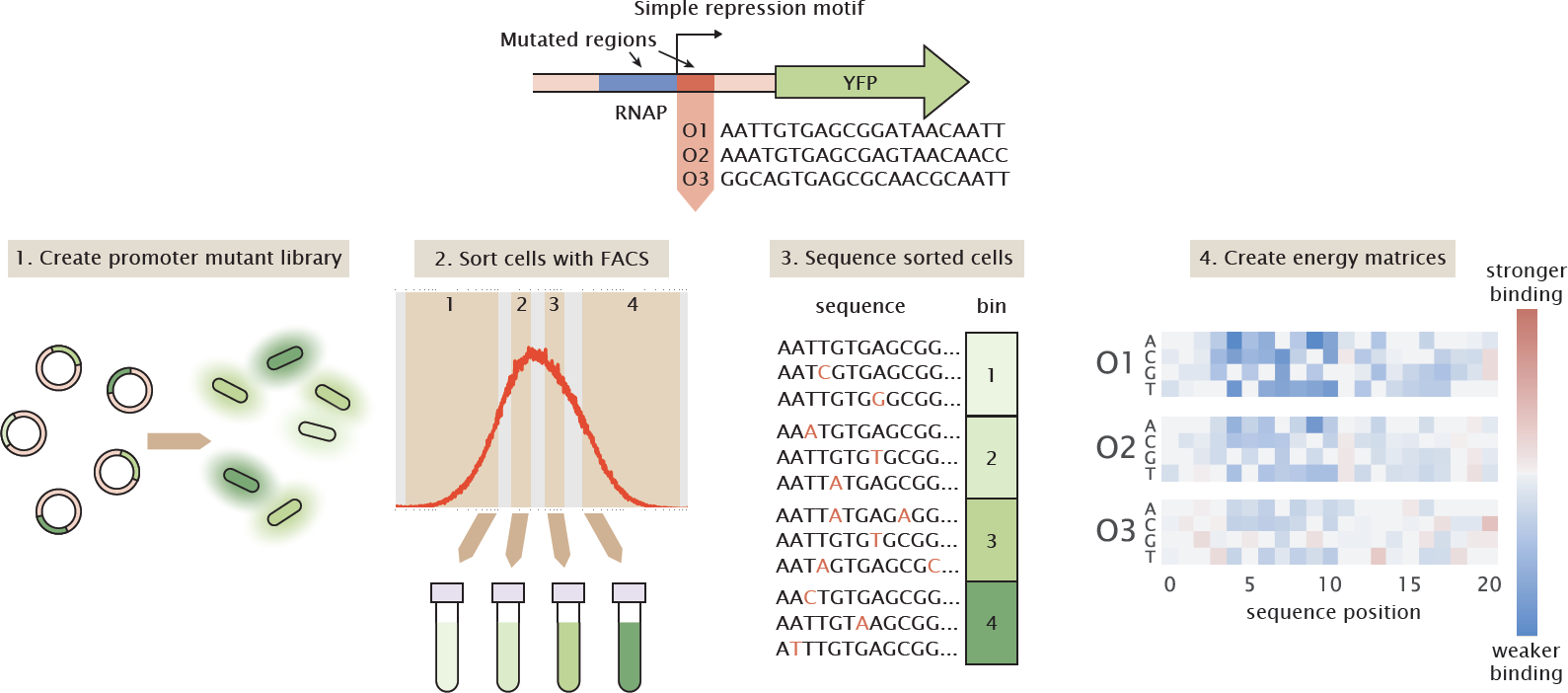
Using Sort-Seq to obtain energy matrices. To begin, we design a simple repression motif in which a repressor binding site is placed immediately downstream of the RNAP site. When RNAP binds, it initiates transcription of the GFP reporter gene. We analyze simple repression constructs using each of the three natural *lac* operators, O1, O2, and O3. Sort-Seq then proceeds as follows. 1. We create a mutant library in which the RNAP and operator sequences are randomly mutated at a rate of approximately 10%, and transform this library into a cell population such that each cell contains a different mutant operator sequence. 2. To measure gene expression, we sort the cell population into bins based on fluorescence level. 3. We then sequence variant promoter sequences within each bin. The bin in which each promoter is found serves as a measure of that promoter’s activity. 4. From this information, we can infer an energy matrix for the repressor binding site indicating which mutations result in a higher or lower binding energy relative to the reference sequence.

As described in Fig 1, to perform Sort-Seq we start by mutating the promoter at a rate of ∼ 10%. Here we mutate both the RNAP binding site and the operator, starting with either O1, O2, or O3 for the operator sequence. While our analysis focuses on the operators themselves, mutating the RNAP site as well aids in model-fitting as described in Appendix B. We place the promoters upstream of a fluorescent reporter gene and create a plasmid library of these constructs. We transform this plasmid library into a population of *E. coli* in which *lacI* and *lacZYA* have been deleted, but *lacI* has been reintroduced to the genome with a synthetic RBS that allows us to precisely control the LacI copy number within the cell, as described in Ref. [29]. We require at least 10^6^ transformants for each plasmid library to ensure sufficient library diversity. Then, we use fluorescence-activated cell sorting (FACS) to sort *E. coli* containing these plasmids into four bins based on their expression levels. We perform high-throughput sequencing on the libraries from each bin. We infer energy matrices that maximize the mutual information between sequence and expression bin (see Appendix B for details). We perform Bayesian parameter estimation using a Markov Chain Monte Carlo algorithm to determine the scaling factor that should be applied to the energy matrix to convert each position into *k*_*B*_*T* energy units. We infer the scaling factor using the same data set that was used to infer the energy matrix, as the ideal scaling factor should maximize the mutual information between promoter sequence and gene expression (see Appendix C for a comparison to other methods for obtaining the scaling factor). At this point, one can compute the expected binding energy of any operator mutant within several mutations of the reference sequence by simply adding together the energy values associated with each base in the operator mutant.

### Choice of reference sequence can alter the repressor’s apparent sequence specificity

One might assume that affinity experiments should reveal the same binding specificity regardless of the set of binding site mutants used in the experiment. To test this possibility, we generated energy matrices using three different reference sequences. A reference sequence refers to the sequence which serves as the “wild-type” for each experiment. For each library, the promoter is mutated relative to its reference sequence. Additionally, when assigning binding energies to an energy matrix, all binding energies are calculated relative to the reference sequence. For our reference sequences we use the three natural *E. coli lac* operators (O1 = AATTGTGAGCGGATAACAATT, O2 = AAATGTGAGCGAGTAACAACC, and O3 = GGCAGTGAGCGCAACGCAATT). For our primary analysis we use energy matrix models. These models assume that each nucleotide position within a binding site contributes independently to the binding energy (see Appendix D for predictions using higher-order models). Each operator has a distinct LacI binding energy, with O1 being the strongest at -15.3 *k*_*B*_*T*, O2 being the second strongest at -13.9 *k*_*B*_*T*, and O3 being the weakest at -9.7 *k*_*B*_*T* [29]. The operator sequences are rather dissimilar to each other, with O2 having 5 mutations relative to O1 and O3 having 8 mutations relative to O1 (and 11 mutations relative to O2). For each library, the average operator sequence has only 2 mutations relative to the reference sequence. As a result, a library generated with O1 as the reference sequence is unlikely to share any mutant sequences with a library generated with O2 or O3 as the reference sequence. Here we assess whether dissimilar mutant libraries generated from different reference sequences produce similar energy matrices and sequence logos from their respective Sort-Seq data sets.

As shown in Fig 2A, the three operators each produce qualitatively similar energy matrices, with the left side of the binding site showing greater sequence dependence than the right side, as evidenced by the larger magnitude of the binding energies assigned to each matrix position. Note that we set the binding energy of the reference sequence to 0 *k*_*B*_*T* for these energy matrices, so that the binding energies assigned to each possible mutation are calculated relative to the reference sequence. For all energy matrices, positions 4-10 show the greatest sequence preference. This preference is reflected in the natural *lac* operator sequences themselves, as the bases from 4-10 are conserved in each of the operators. Notably, the majority of mutations available to O1 incur a penalty to binding energy, while many of the mutations available to O3 enhance the binding energy. This is consistent with the observation that O1 has a strong binding energy while O3 has a weak binding energy.

**Figure 2.**
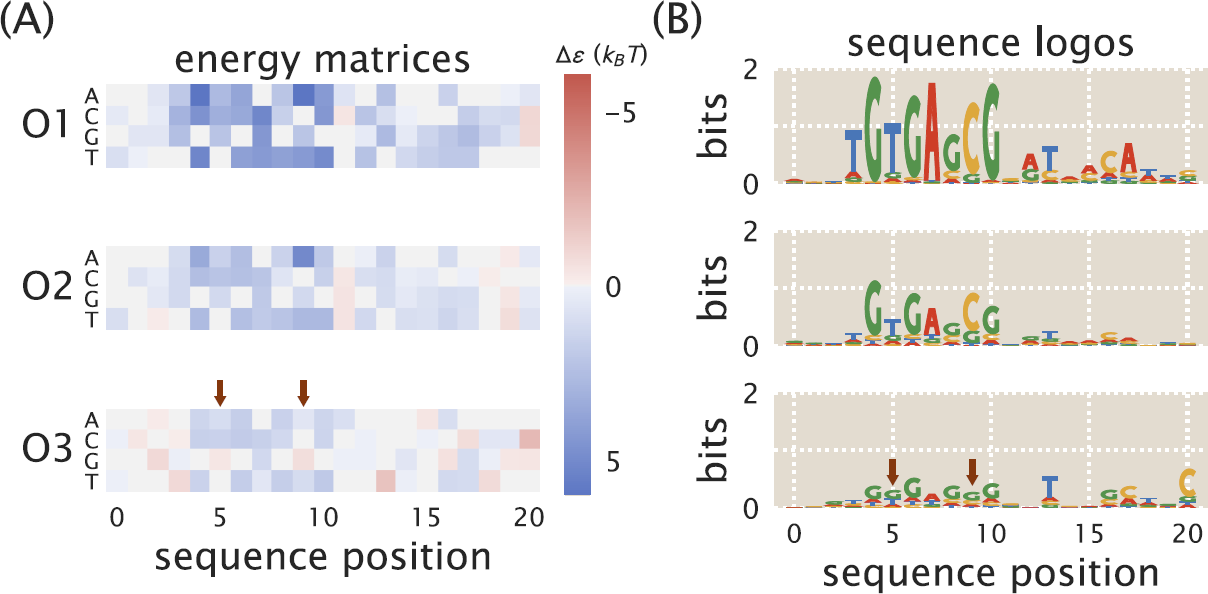
Energy matrices and sequence logos for the natural *lac* operators. A: Energy matrices show how mutations can be expected to affect binding energy. Reference sequences for each energy matrix (either the O1, O2, or O3 sequence) have been set at 0 *k*_*B*_*T* (gray squares), and the energy values at all other positions of the matrix are thus relative to the reference sequence. Red squares represent mutations that create a stronger binding energy than the reference sequence, and blue squares represent mutations that create a weaker binding energy. In columns where multiple squares are gray, this indicates that there is no significant change in binding energy relative to the reference sequence. Positions where preferred bases differ substantially from the O1 matrix are noted with arrows. B: While the energy matrices are qualitatively similar for all three operators, the sequence logos indicate clear differences in the information that can be provided by each operator. The O1 and O2 operators produce similar sequence logos, but the O3 sequence logo incorrectly predicts the preferred binding sequence for LacI. The O3 sequence logo also indicates a much lower information content than for O1 and O2. Positions where preferred bases differ substantially from the O1 sequence logo are noted with arrows.

When the energy matrices are used to produce sequence logos (see Ref. [30]), we see a consistent preference for a slightly asymmetric binding site, reflecting the fact that LacI is known to bind asymmetrically to its operators [31]. Additionally, clear differences arise for the different operators (see Fig 2B). One of the most striking differences is the information content of each sequence logo; as the binding energy of the reference sequence grows weaker, the average information content of each nucleotide position grows smaller. Additionally, while the sequence logos derived from O1 and O2 indicate very similar sequence preferences, the preferred sequence suggested by the O3 sequence logo differs in some prominent positions. In Appendix E we note that weaker binding sites exhibit a greater variation in the quality of their sequence logos; thus it may be that the O3 binding site is simply too weak to provide an informative sequence logo.

### Energy matrix models predict measured energy values

The energy matrices obtained via Sort-Seq should allow us to map sequence to phenotype. The relevant phenotype for simple repression constructs is the degree to which the system is repressed, which can be measured using the fold-change. We define fold-change as the ratio of expression in a repressed system to expression in a system with no repressors, as described by the equation

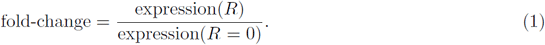

As discussed in further detail elsewhere [28, 29], the fold-change can also be computed using a thermodynamic model given by

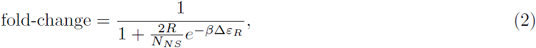

where *R* is the repressor copy number, and the factor of 2 indicates that for the case of LacI, each LacI tetramer has two heads and can essentially be counted as two repressors. *N*_*NS*_ is the number of nonspecific binding sites available in the genome (∼ 4.6 × 10^6^ in *E. coli*) and Δ*ε*_*R*_ is the operator binding energy. We note that this model makes the simplifying assumption that the RNAP binds weakly to the promoter.

In principle, the energy matrix models shown in Fig 2 can be used to predict the binding energy of an operator mutant. To explore the ability of energy matrices to predict the effects of mutations on operator binding strength, we designed a number of mutant operators with 1, 2, or 3 mutations relative to the O1 operator. Experimentally-determined values for the binding energies of these mutants could then be compared against values predicted by our LacI energy matrices.

To obtain experimental values for mutant binding energies we start with chromosomally-integrated simple repression constructs for each mutant, which were incorporated into strains with LacI tetramer copy numbers of *R* = 11 ± 1, 30 ± 10, 62 ± 15, 130 ± 20, 610 ± 80, and 870 ± 170. The error in these copy numbers denotes the standard deviation of at least three Western blot replicates as measured in Ref. [29]. We determined the fold-change by measuring the YFP fluorescence levels of each strain by flow cytometry and substituting them into Eq 1. We determine each mutant’s binding energy, Δ*ε*_*R*_, by performing a single-parameter fit of Eq 2 to the resulting data via nonlinear regression. Fig 3A shows several fold-change values for 1 bp, 2 bp, and 3 bp mutants overlaid with these fitted curves (the remaining fold-change data are shown in Appendix G). To provide a sense of scale for how inaccuracies in binding energy predictions might affect the expected fold-change, the fitted curves are surrounded by a colored region representing Δ*ε*_*R*_ ± 1 *k*_*B*_*T*.

**Figure 3.**
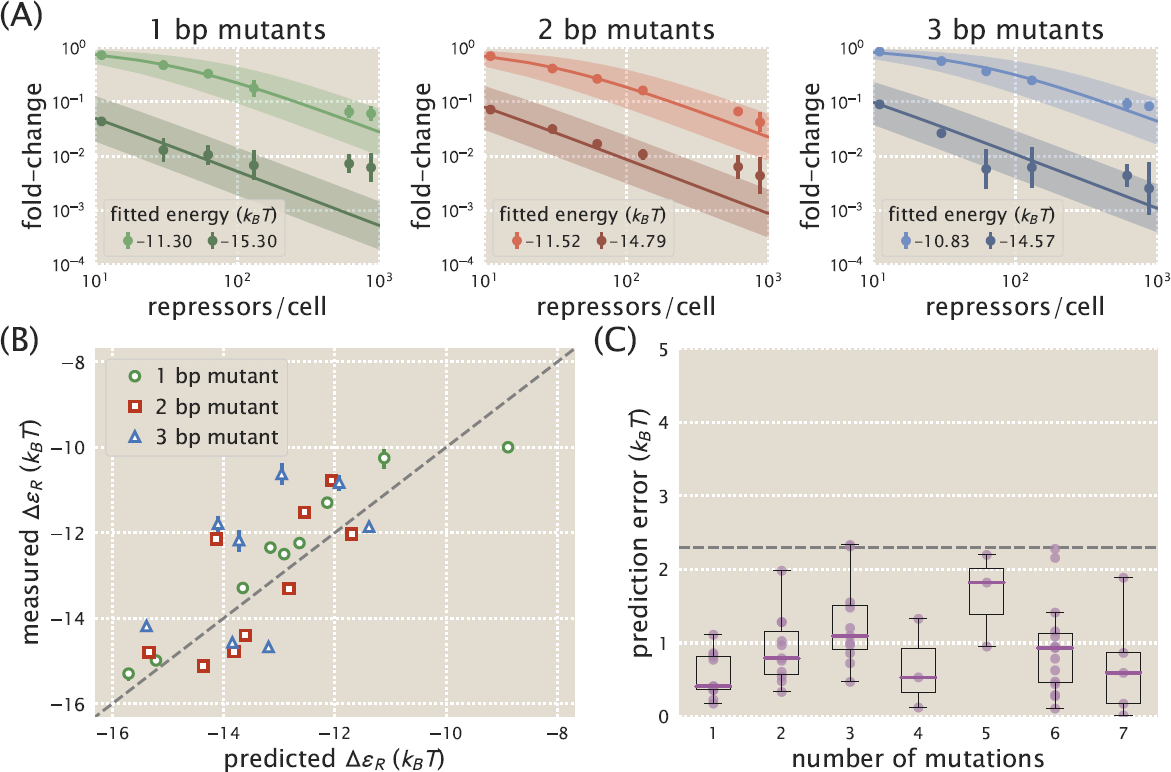
Energy matrix predictions compared to binding energies derived from fold-change data. A: Fold-change data were obtained by flow cytometry for each of the mutant operators by measuring their respective fluorescence levels at multiple LacI copy numbers and normalizing by the fluorescence when *R* = 0. The solid lines in each plot represent a fold-change curve that has been fitted to the data set to obtain a binding energy measurement. The colored region surrounding each fold-change curve indicates the error in fold-change prediction that would result from an error in binding energy prediction of ± 1 *k*_*B*_*T*. Each plot shows data and fits for two operator mutants, one weak and one strong, for 1 bp (left), 2 bp (middle), and 3 bp (right) mutants. All remaining data are shown in Appendix G. Approximately 30 operator mutants were measured in total. We note that lower expression measurements are less accurate than higher expression measurements due to autofluorescence and limitations in the flow cytometer’s ability to measure weak signals. This adversely affects the accuracy of fold-change values for strongly repressed strains. B: The measured binding energy values Δ*ε*_*R*_ (y axis) are plotted against binding energy values predicted from an energy matrix derived from the O1 operator (x axis). While the quality of the binding energy predictions does appear to degrade as the number of mutations relative to O1 is increased, the O1 energy matrix is still able to approximately predict the measured values. C: Binding energies for each mutant were predicted using both the O1 and O2 energy matrices and compared against measured binding energy values. The prediction error, defined as the magnitude of the difference in *k*_*B*_*T* between a predicted binding energy and the corresponding measured binding energy, is plotted here against the number of mutations relative to the reference sequence whose energy matrix was used to make the prediction. Each data point is shown in purple, and box plots representing the data are overlaid to clearly show the median error and variability in error. For sequences with 4 or fewer mutations, the median prediction error is consistently lower than 1.5 *k*_*B*_*T*. The dashed horizontal line represents the point at which the error corresponds to an approximately 10-fold difference in fold-change.

The energy matrices derived from Sort-Seq can be used to predict the value of Δ*ε*_*R*_ associated with a given operator mutant, as discussed in detail in Appendix B. Fig 3B shows how binding energy values measured by fitting to repressor titration data compare to values predicted using energy matrices. For single base pair mutations most predictions perform well and are accurate to within 1 *k*_*B*_*T*, with many predictions differing from the measured values by less than 0.5 *k*_*B*_*T*. Predictions are less accurate for 2 bp or 3 bp mutations, although the majority of these predictions are still within 1.5 *k*_*B*_*T* of the measured value.

The quality of matrix predictions degrades as mutants deviate farther from the wild-type sequence used to generate the energy matrix. To evaluate predictions for a broader range of deviations from the energy matrix, we made predictions from both the O1 energy matrix and the O2 energy matrix. This allowed us to access predictions for operators that are mutated by several base pairs relative to the matrix. In Fig 3C we show how prediction error, defined as the discrepancy in *k*_*B*_*T* between a predicted and measured energy value, varies depending on the number of mutations relative to the wild-type binding site sequence. We find that predictions remain relatively accurate for mutants that differ by up to 4 bp relative to the wild-type sequence, with median deviations of ∼ 1.5 *k*_*B*_*T* or less from the measured binding energy. Other studies have noted that energy matrix models that don’t account for epistatic interactions fail to accurately predict binding energies for mutants with multiple mutations relative to the reference sequence [32, 33]. Thus we find that the relatively low errors depicted in Fig 3C exceed expectations for what a such an energy matrix model can achieve.

We note that energy matrix quality, as measured by the accuracy of its predictions, may be affected by the experimental design. In Appendix E, we assess whether energy matrix quality is affected by the LacI copy number of the background strain, and find that it has little effect on matrix quality. Additionally, we compare predictions made from energy matrices with different reference sequences (i.e. O1, O2, or O3), and find that using O1 as a reference sequence produces the most accurate energy matrices, while using O3 produces energy matrices that are almost entirely non-predictive. In Appendix F, we consider whether better energy matrices are made using libraries in which the entire promoter is mutated or only the operator is mutated. We find that mutating the operator alone can provide more accurate energy matrices, though one must fit energy matrix predictions to binding energy measurements in order to convert these matrices into *k*_*B*_*T* units.

### Designed induction responses

Our predictive energy matrices suggest a promising strategy for addressing the challenge of genetic circuit design, which has typically relied on trial and error to achieve specific outputs [34, 35]. By contrast, previous studies have shown how thermodynamic models can be used to predict gene outputs given a set of inputs [28, 29], which can suggest appropriate inputs to produce a desired output. For example, the key inputs for the fold-change Eq 2 are repressor copy number *R* and repressor-operator binding energy Δ*ε*_*R*_, and one can use Eq 2 to determine a set of *R* and Δ*ε*_*R*_ values that can be used to target a desired fold-change response. Energy matrix predictions can be used to design operator sequences with a particular value of Δ*ε*_*R*_, thereby making it possible to tune genetic circuits and target specific phenotypes. As shown in Fig 3B, mutating an operator by as little as one base pair can provide a broad range of Δ*ε*_*R*_ values that can be predicted accurately.

One particularly useful class of simple genetic circuit, which can be layered with other genetic components to create complex logic [36], is inducible simple repression [37–40]. In such a system, an allosteric repressor can switch between an active form, which binds to an operator with high affinity, and an inactive form, which has a low affinity to the operator. An inducer may bind to the repressor and stabilize the repressor’s inactive form, thereby reducing the probability that the repressor will bind to the operator and increasing the probability that RNAP will bind and initiate transcription. The result is that an inducible system can access a broad range of fold-change values simply by tuning the concentration of inducer. As discussed in Ref. [41], the fold-change of an inducible simple repression circuit can be described by the equation

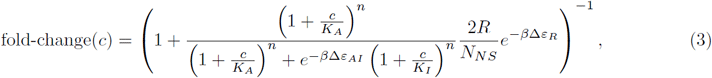

where *c* is the concentration of inducer, *n* is the number of inducer binding sites on the repressor, *K*_*A*_ and *K*_*I*_ are the dissociation constants of the inducer and repressor when the repressor is in its active or inactive state, respectively, and Δ*ε*_*AI*_ is the difference in free energy between the repressor’s active and inactive states. In Ref. [41] we determined that these values are 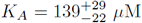, 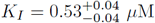, and Δ*ε*_*AI*_ = 4.5 *k*_*B*_*T* for LacI with the inducer IPTG. Where noted, superscripts and subscripts indicate the upper and lower bounds for the 95th percentile of the parameter value distributions. There are *n* = 2 inducer binding sites on each LacI dimer.

We can use these parameter values for the *lac*-based system considered here to explore how tuning the operator-repressor binding energy Δ*ε*_*R*_ can alter the induction response when an effector (i.e. IPTG) is introduced to the system. Importantly, our sequence-energy mapping provides a straightforward avenue for tuning Δ*ε*_*R*_ by altering the binding sequence rather than mutating the repressor itself, which is much more difficult to characterize. We note that an induction response can be described by a number of key phenotypic parameters. The leakiness is the minimum fold-change when no inducer is present, given by fold-change(*c*→0) (Eq S10 in Appendix H). The saturation is the maximum fold-change when inducer is present at saturating concentrations, given by fold-change(*c*→ *∞*) (Eq S11 in Appendix H). The dynamic range is the difference between the saturation and leakiness, and represents the magnitude of the induction response (Eq S13 in Appendix H). The [*EC*_50_] is the inducer concentration at which the fold-change is equal to the midpoint of the induction response (Eq S15 in Appendix H). Full expressions for these parameters are shown in Appendix H. Figs 4A and 4B show how these phenotypic parameters vary with Δ*ε*_*R*_ given the values of *K*_*A*_, *K*_*I*_, and Δ*ε*_*AI*_ listed above and the repressor copy number *R* = 130. We can see that there are inherent trade-offs between phenotypic parameter values. For instance, in this particular system one cannot tune Δ*ε*_*R*_ to obtain a small dynamic range (e.g. a dynamic range of 0.1) while also having an intermediate leakiness value (e.g. a leakiness of 0.4). Rather, one must design an induction response by choosing from the available phenotypes, or else alter the system by tuning additional parameters such as *K*_*A*_ and *K*_*I*_, which requires mutating the protein itself or using a different transcription factor altogether as in Ref. [34].

**Figure 4.**
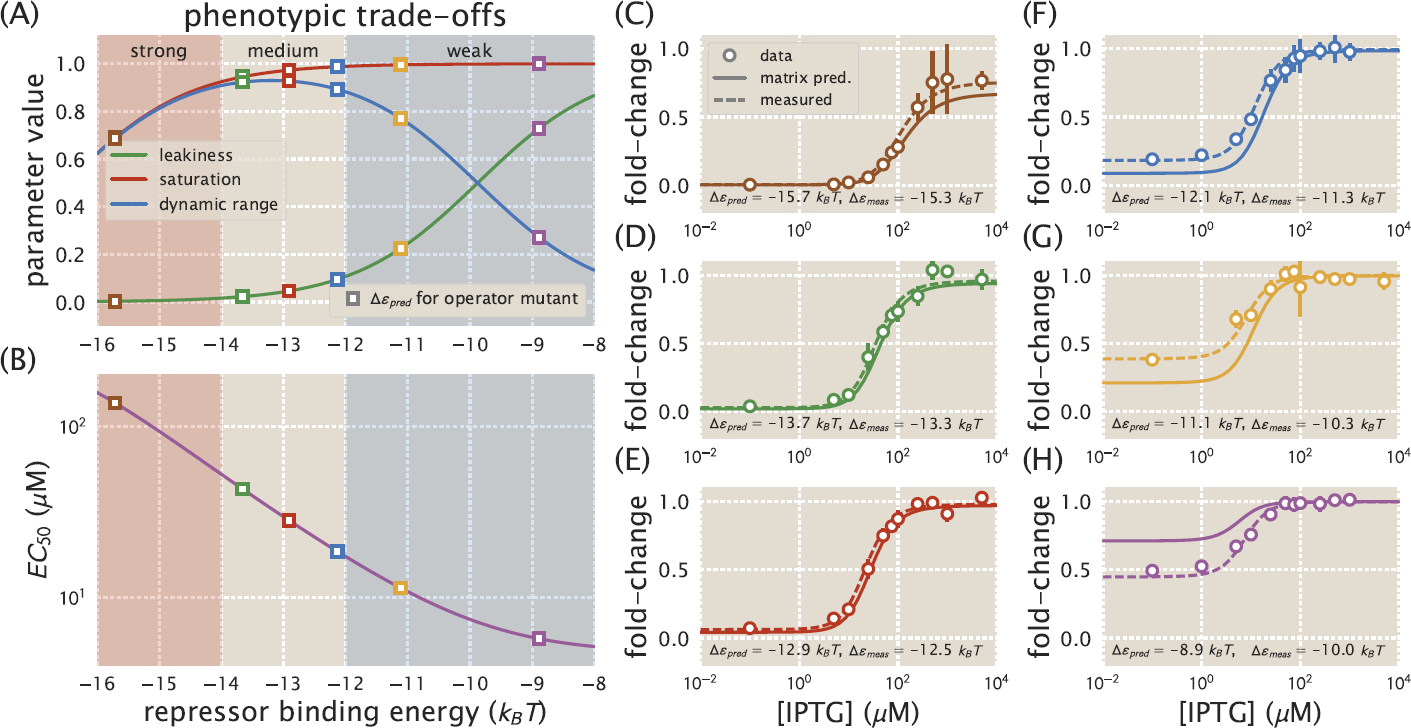
Energy matrix predictions can be used to design phenotypic responses. Phenotypic parameters exhibit trade-offs as Δ*ε*_*R*_ is varied. A: The values of the leakiness, saturation, and dynamic range are plotted as a function of transcription factor binding energy, Δ*ε*_*R*_, for a strain with *R* = 130. Different values of Δ*ε*_*R*_ fall into different binding regimes (strong, medium, or weak) with different phenotypic properties. Several operators were chosen whose predicted binding energies (squares) fall into these different binding regimes. B: The value of the [*EC*_50_] is plotted as a function of Δ*ε*_*R*_ for a strain with *R* = 130. The [*EC*_50_] decreases as the value of Δ*ε*_*R*_ increases. C-H: Operators with different values of Δ*ε*_*R*_ were chosen to have varying induction responses based on the phenotypic trade-offs shown in (A) and (B). The fold-change is shown for each operator as IPTG concentrations are varied. C-E: For operators with stronger binding energies, the data match well with both the predicted theory curves and the theory curves based on measured binding energies. F-H: For operators with weaker binding energies, the data match well with theory curves based on measured binding energies, but do not match as well with predicted theory curves, due to inaccuracies in the energy matrix predictions. For each of these operators, the predicted binding energy Δ*ε*_*pred*_ differs from the measured binding energy Δ*ε*_*meas*_ by ∼ 1 *k*_*B*_*T*.

To show how energy matrices can be used to design specific induction responses, we used the phenotypic trade-offs shown in Figs 4A and 4B to choose values of Δ*ε*_*R*_ that would provide distinct outputs. A strong binding energy lies below Δ*ε*_*R*_ ≈ *-*14 *k*_*B*_*T*, which provides a minimal leakiness level but not full saturation, and gives a high [*EC*_50_] value. A moderate binding energy lies in the range Δ*ε*_*R*_ ≈ *-*14 to *-* 12 *k*_*B*_*T*, maximizing dynamic range and giving an intermediate [*EC*_50_] value. Finally, weak binding energies lie above Δ*ε*_*R*_ ≈ *-*12 *k*_*B*_*T*, which provides a narrower dynamic range and a lower [*EC*_50_] value. We chose six of our single base-pair mutants with predicted binding energies in these ranges. Induction responses for each of these mutants were measured by growing cultures in the presence of varying IPTG concentrations and measuring the fold-change at each concentration. Fig 4C-H shows how the induction data compare against theory curves plotted using Δ*ε*_*R*_ values predicted from the energy matrix. For operators with stronger binding energies, the data match well with the theory curves plotted using predicted binding energies (Fig 4C-E). For operators with weaker binding energies, however, we find that the data do not match as well with the predicted theory curves (Fig 4F-H). Theory curves plotted using the measured binding energy (rather than the predicted binding energy) match well with the data, indicating that the mis-match between the data and the predicted theory curve is due to error in the predicted binding energy.

### Analysis of amino acid-nucleotide interactions

Predictive energy matrices offer a simple way of analyzing direct interactions between amino acids and nucleotides. Mutating individual amino acids in the repressor’s DNA-binding domain and then observing changes in the energy matrix makes it possible to determine how changing the amino acid composition of the DNA-binding domain alters sequence preference. If sequence specificity is altered only for specific base pairs when an amino acid is mutated, this may indicate that the amino acid interacts directly with those base pairs. While it is possible to obtain such information using binding assays [42] or labor-intensive structural biology approaches, Sort-Seq makes it possible to efficiently sample protein-DNA interactions. To analyze the effects of amino acid mutations on sequence specificity, we chose mutations which had previously been found to alter LacI-DNA binding properties without entirely disrupting the repressor’s ability to bind DNA [42, 43]. We performed Sort-Seq using strains containing one of three LacI mutants, Y20I, Q21A, or Q21M, where the first letter indicates the wild-type amino acid, the number indicates the amino acid position, and the last letter indicates the identity of the mutated amino acid.

The energy matrices for each LacI mutant are shown in Fig 5A, along with the wild-type energy matrix for comparison. Sequence logos derived from each energy matrix are shown in Fig 5B. The energy matrices remain remarkably similar to one another. As with the wild-type repressor, for each of the mutant repressors we find that the left half-site of the sequence logo has a stronger sequence preference. For both Y20I and Q21M, the same sequence is preferred in the left half-site as for the wild-type LacI. This contrasts with the results from Ref. [42], in which it was found that Y20I prefers an adenine at sequence position 6, rather than the guanine preferred at this position by the wild-type repressor. As in Ref. [42], we find that an adenine is preferred at sequence position 6 for the Q21A mutant. Additionally, when comparing the left and right half-sites of each energy matrix, we find that for each mutant the preferred sequence is not entirely symmetric. Thus we see that the *lac* repressor’s notable preference for a pseudo-symmetric operator is preserved in each of the mutants we tested.

**Figure 5.**
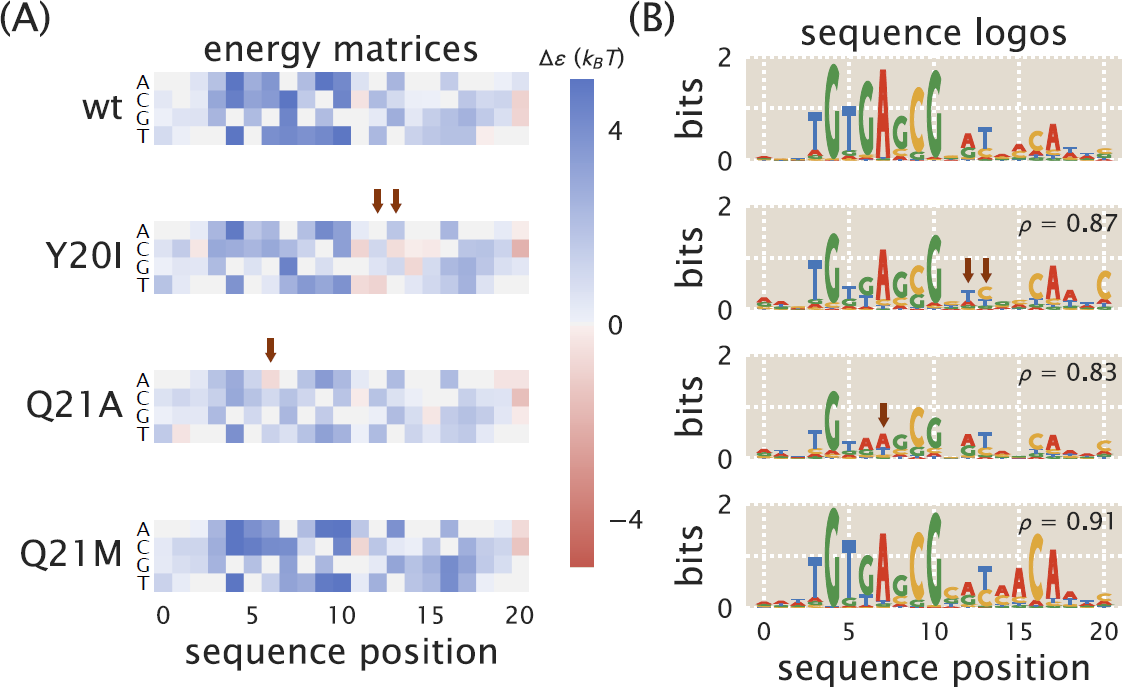
Mutations to LacI DNA-binding domain cause subtle changes to sequence specificity. Mutations were made to residues 20 and 21 of LacI, both of which lie within the DNA-binding domain. The mutations Y20I and Q21A weaken the repressor-operator binding energy, while the mutation Q21M strengthens the binding energy [43]. The sequence preferences of each mutant are represented as A: energy matrices and B: sequence logos. Y20I exhibits minor changes to specificity in low-information regions of the binding site, and Q21A experiences a change to specificity within a high-information region of the binding site (see arrows). Specifically, Q21A prefers A at operator position 6 while the wild-type repressor prefers G at this position. The Pearson’s correlation coefficient *ρ* is noted for each mutant, calculated by comparing the energy matrix values for each mutant to the wild-type energy matrix values. For comparison, replicates of the O1 energy matrix with wild-type LacI all have values of *ρ ≥* 0.93 relative to one another (see Appendix E).

### Binding site context can influence a transcription factor’s binding specificity

In this work we have used the *lac* system to demonstrate how Sort-Seq can be used to map binding site sequence to binding energy, and we used these mappings to rationally design novel genetic circuit elements and identify the effects of amino acid mutations on LacI’s sequence specificity. Importantly, this approach is not specific to the *lac* system and can be applied to any system in which transcription factors alter gene expression by binding to DNA within the promoter region. In Ref. [44] we showed how Sort-Seq could be used alongside mass spectrometry to determine the locations of transcription factor binding sites in a promoter of interest and identify which transcription factors bind to these sites. We generated energy matrices for a number of transcription factors (e.g. RelBE, MarA, PurR, XylR, and others). Here we analyze selected energy matrices from Ref. [44] to show how energy matrices can be used to understand transcriptional activity in promoters with varied architectures beyond simple repression.

One of the questions we wish to answer is to what extent altering the context of a binding site within a regulatory architecture will alter sequence specificity. One hypothesis is that a transcription factor’s preferred binding sequence will remain the same regardless of how its binding site is positioned within the regulatory architecture. However, it is known that factors beyond the core operator sequence, such as flanking sequences and DNA shape, can affect sequence specificity [19, 45, 46]. Additionally, interactions with other proteins may alter the way a transcription factor contacts the DNA, which could affect sequence specificity as well [47]. It is important to know whether a transcription factor’s specificity is sensitive to the context of the binding site within the promoter architecture, as this determines the extent to which an energy matrix can be used to analyze binding sites throughout the genome. Additionally, observing how sequence specificities change with binding site context may alert us to changes in regulatory mechanisms as the operator is moved to different positions in the promoter.

In Ref. [44], we used Sort-Seq to obtain energy matrices and sequence logos for the transcription factors XylR and PurR in the context of the natural promoters for *xylE* and *purT*, respectively. The *xylE* promoter has two XylR binding sites directly adjacent to one another, allowing us to compare these two energy matrices against each other. In this context, we find that XylR appears to act as an activator in tandem with a CRP binding site. Sequence logos for the two XylR binding sites are shown in Fig 6A. The energy matrices and sequence logos for these binding sites have some significant dissimilarities. Dissimilarities are particularly notable at positions 6-8, where the left-hand site prefers “TTT” and the right-hand site prefers “AAA”. In the *xylE* promoter the left-hand XylR site is adjacent to a CRP site, while the right-hand XylR site is adjacent to the RNAP site. The close proximity of these binding sites suggests that there may be direct interactions between proteins, which could alter how each XylR interacts with the DNA, thus altering sequence preferences. The Pearson’s correlation coefficient *ρ* between the two energy matrices is *ρ* = 0.57.

**Figure 6.**
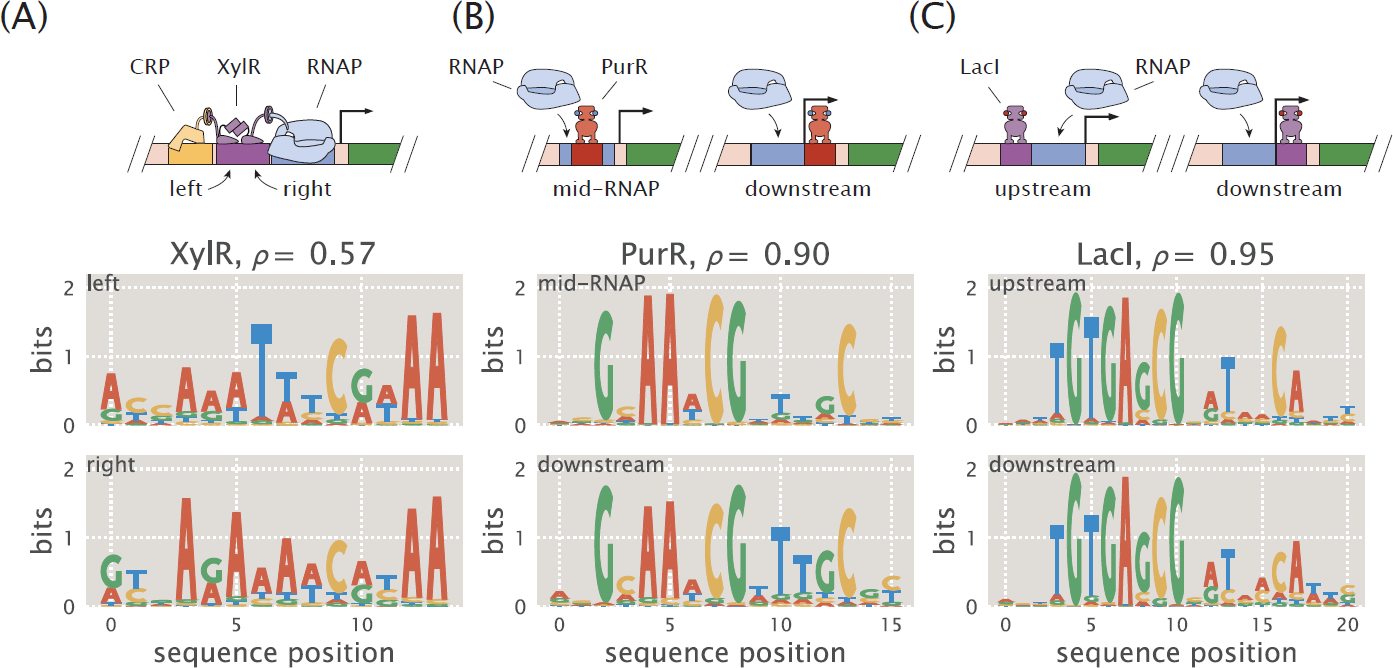
Regulatory context can alter sequence preference. Sequence logos were obtained for the same transcription factors in different regulatory contexts and compared against one another. The Pearson’s correlation coefficient *ρ* between energy matrices is noted for each pair of binding sites. A: Sequence logos are shown for the two adjacent binding sites for the activator XylR in the *xylE* promoter, shown schematically at top. The sequence logos for the two binding sites indicate that they have significantly different sequence preferences. B: Sequence logos are shown for the PurR binding site in the *purT* promoter and a PurR binding site for a synthetic simple repression promoter in which the binding site is positioned differently, shown schematically at top. The sequence logos for the two binding sites indicate nearly identical sequence preferences. C: Sequence logos are shown for a LacI binding site upstream of the RNAP binding site and a LacI binding site downstream of the RNAP. Although regulatory mechanisms differ between these two binding sites, their sequence logos are nearly identical.

In Ref. [44] we find that PurR acts as a repressor in the *purT* promoter, with a single binding site between the -10 and -35 sites. In order to compare the associated energy matrix with a PurR energy matrix from a different regulatory context, here we create a synthetic promoter in which the PurR binding site has been moved directly downstream of the RNAP site. This should continue to be a simple repression architecture in which repressor binding occludes RNAP binding, but the change in operator position may alter the repressor’s interaction with the DNA. Sequence logos for both PurR binding sites are shown in Fig 6B. The two PurR sequence logos are very similar to one another, indicating no significant changes in the interactions between the repressor and the DNA. We calculate the Pearson’s correlation coefficient between the two energy matrices to be *ρ* = 0.90, which is significantly higher than the value calculated for the two XylR energy matrices.

We additionally performed Sort-Seq on a LacI simple repression construct in which the *lac* operator was placed upstream of the RNAP binding site rather than downstream. In Ref. [26] it is shown that LacI binding to an upstream operator still represses, but whereas a downstream operator represses by preventing RNAP from binding, an upstream operator appears to directly contact a bound RNAP and prevent it from escaping the promoter. Moreover, an upstream operator’s binding strength does not directly correspond with the level of repression associated with the promoter. These factors make repression by an upstream *lac* operator an interesting architecture to compare with repression by a downstream *lac* operator. Sequence logos for the upstream and downstream LacI binding sites are shown in Fig 6C. These logos are very similar to one another, despite the fact that the repression mechanisms and protein interactions differ for these two architectures. The Pearson’s correlation coefficient between the two matrices is *ρ* = 0.95.

Because a definitive thermodynamic model was not available for all of the architectures examined in Fig 6, the energy matrices used to make the sequence logos were scaled using a theoretical “average” binding penalty derived from a statistical mechanical analysis of transcriptional regulation (see Appendix C). Appendix A shows the wild-type binding sites that act as reference sequences for the sequence logos.

## Discussion

In this work, we apply quantitative modeling to *in vivo* experimental techniques to analyze interactions between transcription factors and their binding sites under multiple conditions. As an example of how our approach might be used to analyze a transcription factor’s sequence-specific binding energy, we used Sort-Seq to create energy matrices that map DNA sequence to binding energy for the *lac* repressor (Fig 2). We performed this work in the context of a simple repression architecture, which is widespread among bacterial promoters [48] and is frequently used in synthetic biology [40, 49, 50]. We test our model’s predictions against binding energies inferred from fold-change measurements of roughly 30 *lac* operator mutants (Fig 3). These predictions proved to be approximately accurate, even for operators with multiple mutations.

Because we are able to accurately predict operator binding energies, our sequence-energy mappings can be used to design specific regulatory responses, which is of great utility to synthetic biology. We combine energy matrices with a thermodynamic model of inducible simple repression to design induction curves, as demonstrated in Fig 4 [41]. We note that in spite of the overall success of our predictions, there remain some predictions that are significantly different from the measured values (see the outliers in Fig 3C). Such inaccuracies are particularly problematic when using energy matrices for design applications, as discrepancies between a system’s expected and actual response may render a designed system unsuitable for its intended application. We can see examples of this in Fig 4F-H, where inaccuracies in binding energy predictions are reflected in the predicted titration curves. The prediction curves corresponding to operators with weaker binding energies do not accurately describe the data, with the data exhibiting higher or lower leakiness values than was predicted. If the leakiness is a vital parameter in the designed system, then such a mis-match could cause the system to fail.

We also explore how sequence specificity is altered when transcription factor amino acids are mutated. To do this, we repeat our Sort-Seq experiments in bacterial strains expressing LacI mutants in which the DNA-binding domain has been altered (Fig 5). Because all nucleotides in the binding site are mutated with some frequency in Sort-Seq experiments, we are able to identify changes in specificity throughout the entire binding site. Other methods for analyzing the sequence preference of transcription factor mutants tend to be more laborious and less fine-grained, often focusing on a small set of nucleotides within the binding site. These include binding experiments between DNA mutants and protein mutants [42], gene expression experiments using chimeric transcription factor proteins [51], and comparative genomics [52].

We further explore how regulatory context alters sequence specificity. We generate sequence logos from energy matrices obtained for the transcription factors XylR, PurR, and LacI in different regulatory contexts, as shown in Fig 6. We find that the two adjacent XylR binding sites exhibit significantly different binding specificities, possibly due to interactions between transcription factors. In contrast, the simple repression constructs analyzed for PurR and LacI have nearly identical sequence specificities. By itself, our method is unable to determine the causes of context-dependent changes in sequence specificity, though it is known that DNA shape or binding to cofactors can alter a transcription factor’s specificity [45–47]. Rather, our approach can be used to determine whether a given binding site’s sequence preferences diverge from the “standard” sequence specificity for the relevant transcription factor, and further experiments (such as SELEX-seq in the presence of a transcription factor and possible cofactors [47]) can be performed to determine the cause of the change in sequence specificity.

A major advantage of our *in vivo* approach is that it allows us to analyze transcription factors in their natural context, in the presence of interacting proteins, small molecules, and DNA shape effects. This is especially important when analyzing regulatory regions that have not been previously annotated, as was the case for the XylR and PurR matrices obtained in Ref. [44]. However, a clear advantage of *in vitro* approaches is that they can accurately measure low-affinity binding sites [12, 13, 15]. When using our *in vivo* approach, weaker reference sequences produce energy matrices with variable quality and are more likely to make poor predictions (see Appendix E). However, accuracy may be improved by investigating ways to reduce the experimental noise associated with *in vivo* systems, for instance by incorporating promoter constructs as single copies in the chromosome rather than multiple copies on plasmid, for example using the “landing pad” technique described in Ref. [53].

This work provides a foundation for further studies that would benefit from sequence-energy mappings. For example, our analysis of three LacI amino acid mutants could be expanded to include a full array of DNA-binding mutants, which would allow one to make inferences regarding repressor-operator coevolution. Additionally, while we make extensive use of LacI in the present work, similar analyses could be performed with any transcription factor, making it possible to improve upon the genomically-inferred sequence logos presently available for many transcription factors. Further, for cases in which it is known that sequence specificity is affected by DNA shape, flanking sequences, cofactor binding, or other factors outside of the operator binding sequence, our approach can be used to obtain a finely-detailed map of the effects on sequence specificity. Finally, we note that one of the primary strengths of our approach is that it can be used to elucidate the transcriptional regulation of a gene with a previously-unknown regulatory architecture. As shown in Ref. [44], Sort-Seq can be combined with mass spectrometry to identify transcription factor binding sites and those sites’ regulatory roles for any gene of interest. Here we show that data sets obtained in this manner can also be used to map sequence to binding energy, thus showing that a single experiment can be used to characterize multiple aspects of a previously-unannotated regulatory sequence. Futhermore, our approach does not rely specifically on the Sort-Seq technique used here, but can be adapted to multiple experimental designs, such as RNA-seq based MPRAs that have been demonstrated in multiple model systems [7, 54–56]. Over time, we envision incorporating high-throughput synthesis and analysis techniques to adapt our approach for genome-wide studies in both prokaryotes and eukaryotes.

## Methods

### Sort-Seq libraries

To generate promoter libraries for Sort-Seq, mutagenized oligonucleotide pools were purchased from Integrated DNA Technologies (Coralville, IA). These consisted of single-stranded DNA containing the *lacUV5* promoter and LacI operator plus 20 bp on each end for PCR amplification and Gibson Assembly. Either both the *lacUV5* promoter and LacI binding site or only the LacI binding site was mutated with a ten percent mutation rate per nucleotide. These oligonucleotides were amplified by PCR and inserted back into the pUA66-operator-GFP construct using Gibson Assembly. To achieve high transformation efficiency, reaction buffer components from the Gibson Assembly reaction were removed by drop dialysis for 90 minutes and cells were transformed by electroporation of freshly prepared cells. Following an initial outgrowth in SOC media, cells were diluted with 50 mL LB media and grown overnight under kanamycin selection. Transformation typically yielded 10^6^ - 10^7^ transformants as assessed by plating 100 *μ*L of cells diluted 1:10^4^ onto an LB plate containing kanamycin and counting the resulting colonies.

### DNA Constructs for fold-change measurements of mutant operators

Simple repression motifs used in fold-change measurements were adapted from those in Garcia *et al.* [29]. Briefly, a simple repression construct with the O1 operator sequence was cloned into a pZS25 plasmid background directly downstream of a *lacUV5* promoter, driving expression of a YFP gene when the operator is not bound by LacI. This plasmid contains a kanamycin resistance gene for selection. Mutant LacI operator constructs (listed in Table 1) were generated by PCR amplification of the *lacUV5* O1-YFP plasmid using primers containing the point mutations as well as sufficient overlap for re-circularizing the amplified DNA by one-piece Gibson Assembly.

**Table 1.**
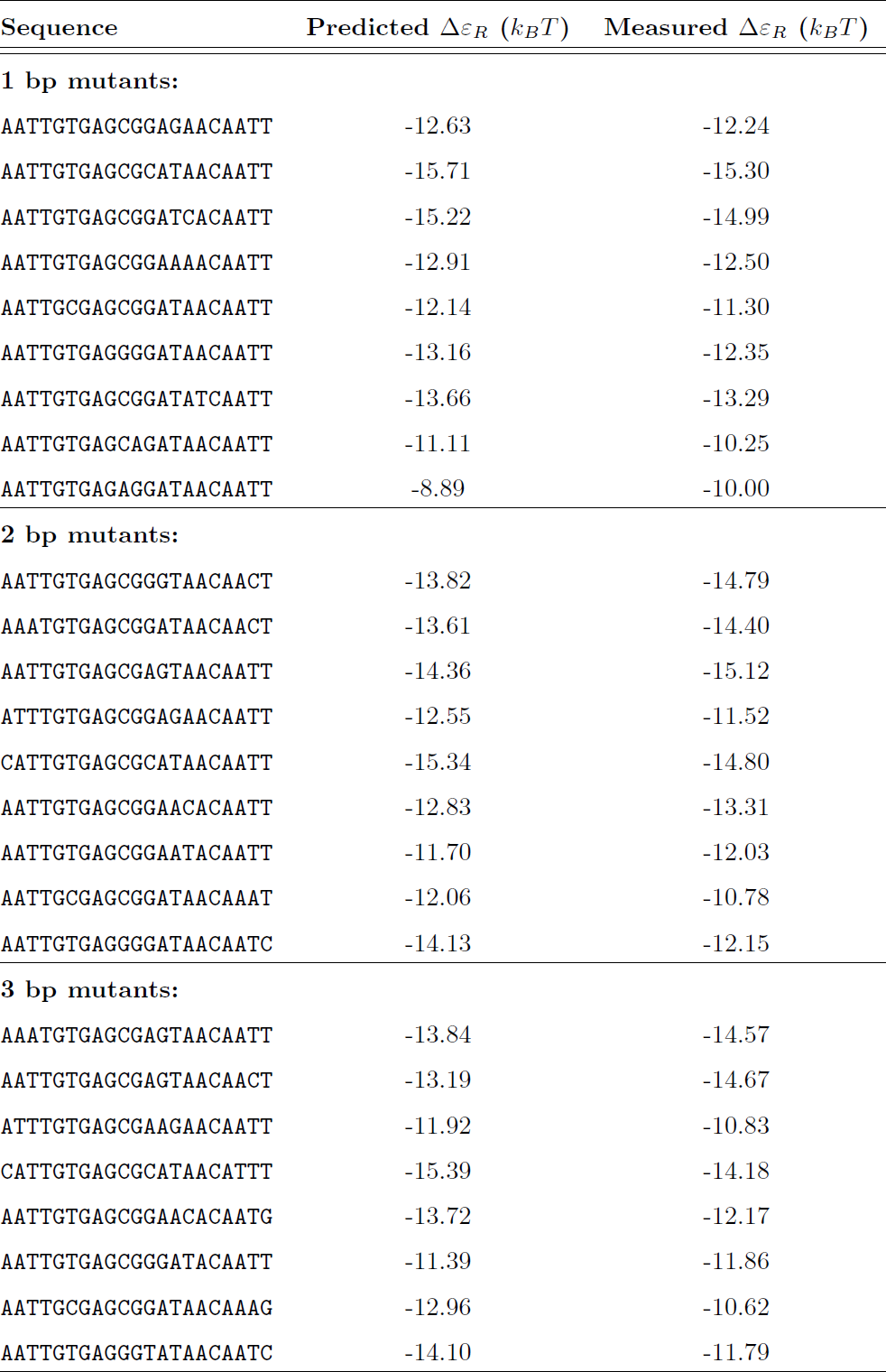
Mutant operator sequences. The listed operator sequences were used to evaluate energy matrix predictions. They are mutated relative to the O1 *lac* operator. The predicted binding energy was generated using the matrix with an O1 reference sequence with *R* = 130 LacI tetramers in the background strain.

A second construct was generated to express LacI at a specified copy number. Specifically, *lacI* was cloned into a pZS3*1 background that provides constitutive expression of LacI from a P_LtetO*-*1_ promoter [57]. This plasmid contains a chloramphenicol resistance gene for selection. The LacI copy number is controlled by mutating the ribosomal binding site (RBS) for the *lacI* gene as described in [58] using site-directed mutagenesis (Quickchange II; Stratagene, San Diego, CA) and further detailed in [29]. Here, we mutated the RBS such that it would produce a LacI copy number of ∼ 130 tetramers once the construct had been integrated into the chromosome.

Once the plasmids had been generated, the promoter and *lacI* constructs were each amplified by PCR and integrated into the chromosome by lambda-red recombineering using the pSIM6 expression plasmid [59]. The promoter construct and YFP gene were inserted into the *galK* locus in the *E. coli* genome and the *lacI* construct was inserted into the *ybcN* locus.

### Construction of LacI Amino Acid Mutants

As previously mentioned, wild-type *lacI* was cloned into a pZS3*1 background providing constitutive expression of LacI, with the LacI copy number mediated by a mutated RBS. We used the RBS corresponding to a LacI tetramer copy number of ∼ 130 for each mutant. To create DNA-binding mutants for LacI we used site-directed mutagenesis (Quickchange II; Stratagene, San Diego, CA) using the mutagenesis primers listed in Table 2. We mutated the amino acid Y to I at position 20 and Q to A or M at position 21. We chose these mutations based on data from previous studies [42, 43], though we note that our amino acid numbering system is shifted by +3 relative to the mutants in these previous studies since we use a slightly different version of *lacI*. As with the wild-type *lacI*, we integrate the mutants into the genome at the *ybcN* locus by lambda-red recombineering using the pSIM6 expression plasmid.

**Table 2.**
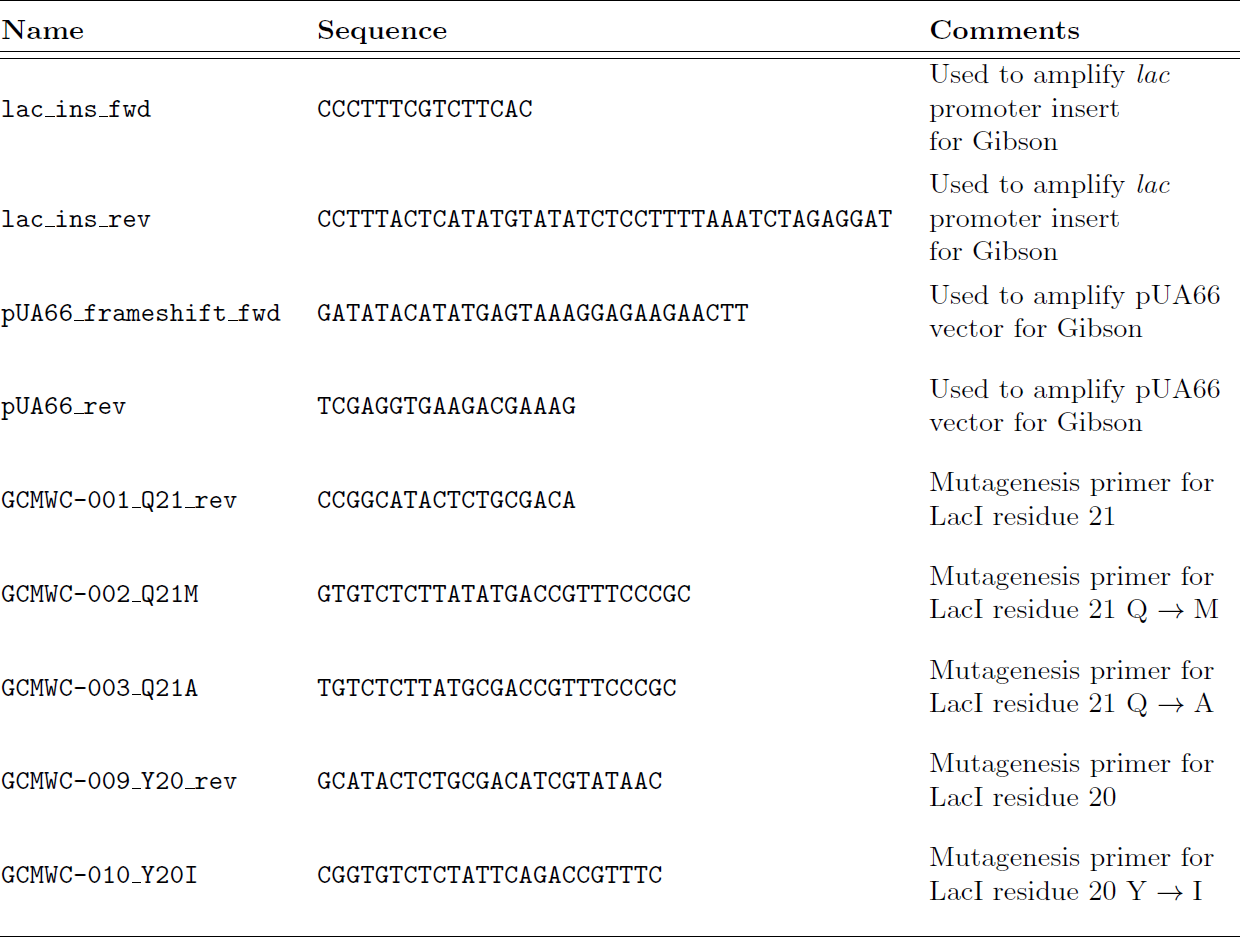
Primers used in this work. The listed primer sequences were used to generate plasmids for Sort-Seq experiments or for use in creating strains with mutated operators or LacI.

### Bacterial Strains

*E. coli* strains used in this work were derived from K12 MG1655. To generate strains with different LacI copy number, the *lacI* constructs were integrated into a strain that additionally has the entire *lacI* and *lacZYA* operons removed from the chromosome. These constructs were integrated at the *ybcN* chromosomal location. This resulted in strains containing mean LacI tetramer copy numbers of *R* = 11 ± 2, 30 ± 10, 62 ± 15, 130 ± 20, 610 ± 80, and 870 ± 170, where the error denotes the standard deviation of at least three replicates as measured by quantitative western blots in Ref. [29].

For Sort-Seq experiments, plasmid promoter libraries were constructed as described below and then transformed into the strains with *R* = 30, 62, 130 or 610. For fold-change measurements, each O1 operator mutant was integrated into strains containing each of the listed LacI copy numbers. These simple repression constructs were chromosomally integrated at the *galK* chromosomal location via lambda red recombineering. Generation of the final strains containing a simple repression motif and a specific LacI copy number was achieved by P1 transduction. For each LacI titration experiment, we also generated a strain in which the operator-YFP construct had been integrated, but the *lacI* and *lacZYA* operons had been removed entirely. This provided us with a fluorescence expression measurement corresponding to *R* = 0, which is necessary for calculation of fold-change.

### Sort-Seq fluorescence sorting

For each Sort-Seq experiment, cells were grown to saturation in lysogeny broth (LB) and then diluted 1:10,000 into minimal M9 + 0.5% glucose for overnight growth. Once these cultures reached an OD of 0.2-0.3 the cells were washed three times with PBS by centrifugation at 4000 rpm for 10 minutes at 4^*°*^C. They were then diluted two-fold with PBS to reach an approximate OD of 0.1-0.15. These cells were then passed through a 40 *μ*m cell strainer to eliminate any large clumps of cells.

A Beckman Coulter MoFlo XDP cell sorter was used to obtain initial fluorescence histograms of 500,000 events per library in the FL1 fluorescence channel with a PMT voltage of 800 V and a gain of 10. The histograms were used to set four binning gates that each covered ∼ 15% of the histogram. 500,000 cells were collected into each of the four bins. Finally, sorted cells were regrown overnight in 10 mL of LB media, under kanamycin selection.

### Sort-Seq sequencing and data analysis

Overnight cultures from each sorted bin were miniprepped (Qiagen, Germany), and PCR was used to amplify the mutated region from each plasmid for Illumina sequencing. The primers contained Illumina adapter sequences as well as barcode sequences that were unique to each fluorescence bin, enabling pooling of the sorted samples. Sequencing was performed by either the Millard and Muriel Jacobs Genetics and Genomics Laboratory at Caltech or NGX Bio (San Fransisco, CA). Single-end 100bp or paired-end 150bp flow cells were used, with about 500,000 non-unique sequences collected per library bin. After performing a quality check and filtering for sequences whose PHRED score was greater than 20 for each base pair, the total number of useful reads per bin was approximately 300,000 to 500,000 per million reads requested. Energy weight matrices for binding by LacI and RNAP were inferred using Bayesian parameter estimation with a error-model-averaged likelihood as previously described [27, 60] and further detailed in S1 Appendix.

### Fold-change measurements by flow cytometry

Fold-change measurements were collected as previously described in Ref. [41] on a MACSquant Analyzer 10 Flow Cytometer (Miltenyi Biotec, Germany). Briefly, YFP fluorescence measurements were collected using 488nm laser excitation, with a 525/50 nm emission filter. Settings in the instrument panel for the laser were as follows: trigger on FSC (linear, 423V), SSC (linear, 537 V), and B1 laser (hlog, 790V). Before each experiment the MACSquant was calibrated using MACSQuant Calibration Beads (Miltenyi Biotec, CAT NO. 130-093-607). Cells were grown to OD 0.2-0.3 and then diluted tenfold into ice-cold minimal M9 + 0.5% glucose. Cells were then automatically sampled from a 96-well plate kept at approximately 4^*°*^ - 10^*°*^C using a MACS Chill 96 Rack (Miltenyi Biotec, CAT NO. 130-094-459) at a flow rate of 2,000 - 6,000 measurements per second.

For those measurements that were taken for IPTG induction curves, cells were grown as above with the addition of an appropriate concentration of IPTG (Isopropyl *β*-D-1 thiogalactopyranoside Dioxane Free, Research Products International). For each IPTG concentration, a stock of 100-fold concentrated IPTG in double distilled water was prepared and partitioned into 100 *μ*L aliquots. The same parent stock was used for all induction experiments described in this work.

The fold-change in gene expression was calculated by taking the ratio of the mean YFP expression of the population of cells in the presence of LacI to that in the absence of LacI. Since the measured fluorescence intensity of each cell also includes autofluorescence which is present even in the absence of YFP, we account for this background by computing the fold change as

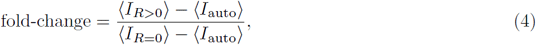

where ⟨*I*_*R>*0_⟩ is the average cell YFP intensity in the presence of repressor, ⟨*I*_*R*=0_⟩ is the average cell YFP intensity in the absence of repressor, and *I*_auto_ is the average cell autofluorescence intensity as determined by measuring the fluorescence of cells in which *R* = 0 and there is no fluorescent reporter.

### Data curation

All data was collected, stored, and preserved using the Git version control software in combination with off-site storage and hosting website GitHub. Code used to generate all figures and perform processing and analyses is available on the GitHub repository (https://www.github.com/rpgroup-pboc/seq_mapping). Inferred model parameters for each energy weight matrix are also available here. Raw flow cytometry data files (.fcs and.csv) files were stored on-site under redundant storage. Due to size limitations, these files are available upon request. Sequencing data is available through the NCBI website under accession number SRP146291.

## Acknowledgements

Access to the Miltenyi Biotec MACSquant Analyzer 10 Flow Cytometer was graciously provided by the Pamela Bjöorkman lab at Caltech. Access to the Beckman-Coulter MoFlo XDP cell sorter was provided by the Tirrell lab, with ongoing technical support from Tirrell lab members Seth Lieblich and Bradley Silverman. Manuel Razo and Griffin Chure from the Phillips lab assisted with constructing the LacI amino acid mutants. Suzannah Beeler from the Phillips lab provided helpful comments on the manuscript draft. This work was supported by the NSF GFRP, the National Institutes of Health DP1 OD000217 (Director’s Pioneer Award), and 1R35 GM118043-01 (MIRA).

## Supporting Information for Mapping operator sequence to transcription factor binding energy *in vivo*

### A Sequences used in this work

**Figure S1.**
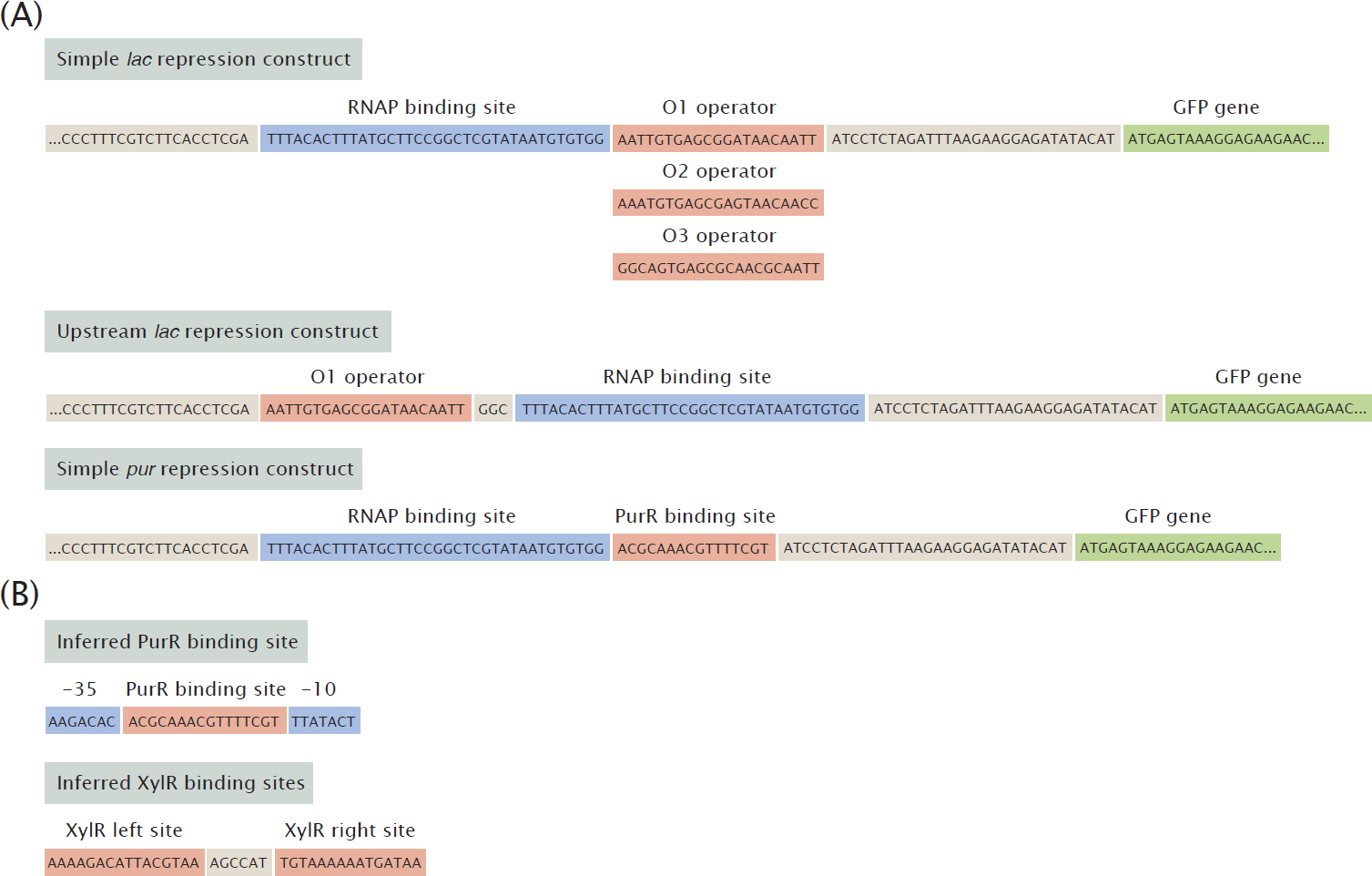
List of wild-type reporter constructs. (A) Wild-type versions of reporter constructs that were used either for Sort-Seq (all) or for measuring operator mutant binding energies (simple *lac* repression). (B) Wild-type versions of sequences that were inferred for PurR and XylR in Ref. [44].

### B Bayesian Inference of Energy Matrix Models

We use Sort-Seq data to generate energy matrices that map sequence to binding energy. As discussed in Refs. [27, 61], one can infer these energy matrices by Bayesian parameter estimation using the observation that for large data sets,

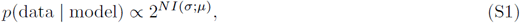

where *N* is the number of data points and *I*(*Σ*; *μ*) represents the mutual information between the promoter sequence *Σ* and the fluorescence bin *μ*. Using a method discussed in detail in Refs. [27, 62], we use a Markov Chain Monte Carlo (MCMC) algorithm to infer a set of energy values (in arbitrary units) for each energy matrix position that maximizes the mutual information between binding site sequence and fluorescence bin. This inference is performed using the MPAthic software package [63].

In order to convert energy matrices into absolute energy units (such as the *k*_*B*_*T* units used in this work), one must obtain a scaling factor that can be applied to the matrix. To obtain this scaling factor, we first observe that energy matrices derived from Sort-Seq can be used to predict the binding energy associated with a given operator mutant (Δ*ε*_*R*_) using the linear equation

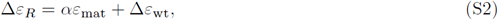

where *ε*_mat_ is the energy value obtained by summing the matrix elements associated with a sequence, *α* is a scaling factor that converts the matrix values into *k*_*B*_*T* units, and Δ*ε*_wt_ is the binding energy associated with the reference sequence. The values of the matrix positions associated with the reference sequence are fixed at 0 *k*_*B*_*T*, so that *ε*_mat_ = 0 for the reference sequence. Thus, *αε*_mat_ can be interpreted as the change in binding energy relative to the reference sequence caused by the specific mutations to the sequence. The value of *α* can be determined in a number of ways (as discussed further in Appendix C), but the method employed in the main text is to use Bayesian parameter estimation by MCMC. The advantage of this method is that if a thermodynamic model for the promoter is known, one can use the Sort-Seq data to infer the value of *α* without having to perform any additional experiments. Here we describe in detail how MCMC is used to infer a value for *α*.

If the energy matrix is properly converted into *k*_*B*_*T* units, then one can use energy matrix predictions, along with a thermodynamic model for gene expression, to discern which fluorescence bin a given promoter sequence should have fallen into. We discuss above how one can infer the energy matrix parameters by maximizing the mutual information between sequence and expression bin. Similarly, we can obtain an estimate for *α* by finding the value of *α* that maximizes the mutual information between the Sort-Seq data and the expression predictions from the matrix and thermodynamic model. For the thermodynamic model, we begin with the expression for *p*_*bound*_ for a simple repression system,

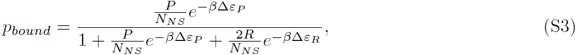

where *P* is the number of RNAP molecules in the system, *N*_*NS*_ is the number of nonspecific binding sites available in the system (i.e. the length of the genome), *R* is the number of repressors in the system, Δ*ε*_*P*_ is the binding energy of RNAP to its binding site, and Δ*ε*_*R*_ is the binding energy of the repressor to its binding site. We can rearrange this equation to make it easier to work with. First, we divide the top and bottom by the numerator, giving us

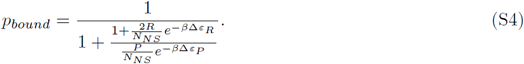

Importantly, in order to evaluate the mutual information between Δ*ε*_*R*_ and *p*_*bound*_, it is not necessary to adhere to the full expression for *p*_*bound*_. Rather, we can manipulate the expression in ways that make it easier for us to work with, provided that the mutual information between Δ*ε*_*R*_ and *p*_*bound*_ is preserved. As noted in [60], the mutual information is preserved provided that any manipulations to the expression do not disrupt the rank ordering of an expression’s values as the value of Δ*ε*_*R*_ is varied. We note that the term 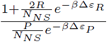 has the same rank ordering as the full expression for *p*_*bound*_. Furthermore, taking the log of this term will also not affect the rank ordering, and it will make the calculation simpler, so we take the log to get an expression which we will refer to as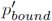, giving us

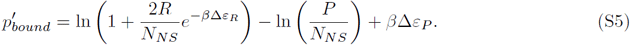

We observe that the constant ln 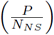 also does not affect rank ordering, so we can drop this term. Additionally, we recall that Δ*ε*_*R*_ = *αE*_*mat,R*_ + Δ*ε*_*wt,R*_. Likewise, we can say that Δ*ε*_*P*_ = *ΓE*_*mat,P*_ + Δ*ε*_*wt,P*_, where *Γ* is the scaling factor for the RNAP matrix. As before, we can drop the constant Δ*ε*_*wt,P*_ as it will not affect rank ordering. This leaves us with the expression

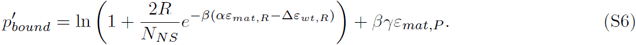

With this expression in hand we can sample values of *Γ* and *α* to identify values that maximize the mutual information between 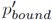 and the expression bin which a particular sequence was sorted into during Sort-Seq. Note that while the rest of the discussion will focus on *α*, a value for *Γ* comes out of this analysis as well.

The mutual information surface is very rough, with many peaks, so we need to use a method which can avoid getting stuck in local maxima. We use a parallel tempering MCMC algorithm to achieve this [64]. The parallel tempering MCMC algorithm works by randomly sampling possible values for *α* and rejecting the value with some probability if it does not increase the mutual information relative to the previous sampled value of *α*. In this respect it is similar to a “standard” MCMC algorithm. By contrast with a standard MCMC algorithm, a parallel tempering algorithm runs multiple chains at once at different temperatures. In our case, we use 10 different temperatures ranging from *β* = 0.02 to *β* = 4 on a log scale, where *β* = 1*/k*_*B*_*T*. Periodically throughout the MCMC run, the current *α* values from different temperature chains will swap. This allows the algorithm to sample *α* values at different levels of precision. Specifically, the high temperature chains will explore widely and not get stuck in local minima, while the low temperature chains will then carefully explore the peak that was found by the high temperature chain. The output is a distribution of values, and we take the median of this distribution to obtain our estimate for *α*.

### C Alternate Methods for Obtaining Energy Matrix Scaling Factor

As discussed in Appendix B, in order to convert an energy matrix into *k*_*B*_*T* units one must infer an appropriate scaling factor *α*. In the main text we primarily use Bayesian parameter estimation by MCMC to infer this factor, but other methods can be used as well. Here we discuss two alternative methods: least squares regression to measured binding energy values, and calibrating to a theoretical mutation parameter. In this Appendix we will discuss the strengths and weaknesses of each method and compare predictions using these methods to predictions using MCMC.

#### Fitting by Least Squares Regression to Measured Binding Energy Values

To obtain a value for *α* using least squares regression, we first define a least-squares function *f* (*α*) as

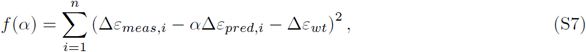

where Δ*ε*_*meas*_ is the measured binding energy for an operator mutant, Δ*ε*_*pred*_ is the corresponding binding energy prediction from our unscaled energy matrix, and Δ*ε*_*wt*_ is the binding energy of the reference sequence used to generate the matrix. To determine the best-fit value of *α*, we identify the value of *α* that minimizes the function. We perform this fit using measurements from the nine single base pair mutants used in this work.

#### Fitting to the Average Energy per Mutation

In many cases, we will not have thermodynamic models available to use for inferring scaling factors by fitting or Bayesian inference. This raises the question of whether it is possible to estimate the scaling factor by other means, for example by determining some average binding penalty incurred by making a mutation to a binding site. To explore how we might think about such an average binding penalty, we consider the effects of mutations away from the lowest-energy binding sequence for LacI (Figure S2). As shown in Figure S2A, a wide range of binding energies are available to binding site mutants. The distribution of binding penalties of single base-pair mutations to this binding site is shown in Figure S2B. The distribution is fairly broad, yet we find that the mean predicted binding energy for binding site mutants, as shown in Figure S2C, is strongly related to the mean binding penalty of a single base pair mutation. Specifically, the slope of the predicted energy versus the number of mutations is approximately equal to the mean binding penalty of a single mutation. This tells us that the average energy per mutation is a meaningful metric that provides information about the general behavior of a transcription factor binding site.

Next we need to determine how one would estimate the average energy per mutation for an energy matrix that has not already been converted into absolute energy units. We turn to Ref. [65] in which they make an estimate for the average energy penalty, *ε*_*mut*_, of a single base pair mutation relative to the minimum-energy sequence. We can use this estimate to infer a value for *α* when no thermodynamic model is available to perform a fit for *α*. We note that unlike the other methods for obtaining *α*, this method does not rely on expression information from the promoter of interest and thus is best interpreted as a “rough guess.”

To begin this estimate, we assume a minimal organism in which there is a single transcription factor with a copy number of 1, and this transcription factor regulates gene expression by binding to a single minimum-energy operator, which has an energy of Δ*ε*_*min*_. The remaining sequence in this minimal genome is mostly random, but it includes a number of weaker binding sites for the transcription factor such that all possible single base-pair mutations to the binding site are represented. From a statistical mechanics perspective, in order for the transcription factor to bind reliably to the minimum-energy operator, the operator’s statistical weight (given by 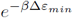) must outweigh the total statistical weight of all possible single base-pair binding site mutants (given by =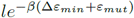), where *l* is the length of the binding site in base pairs. This gives us

**Figure S2.**
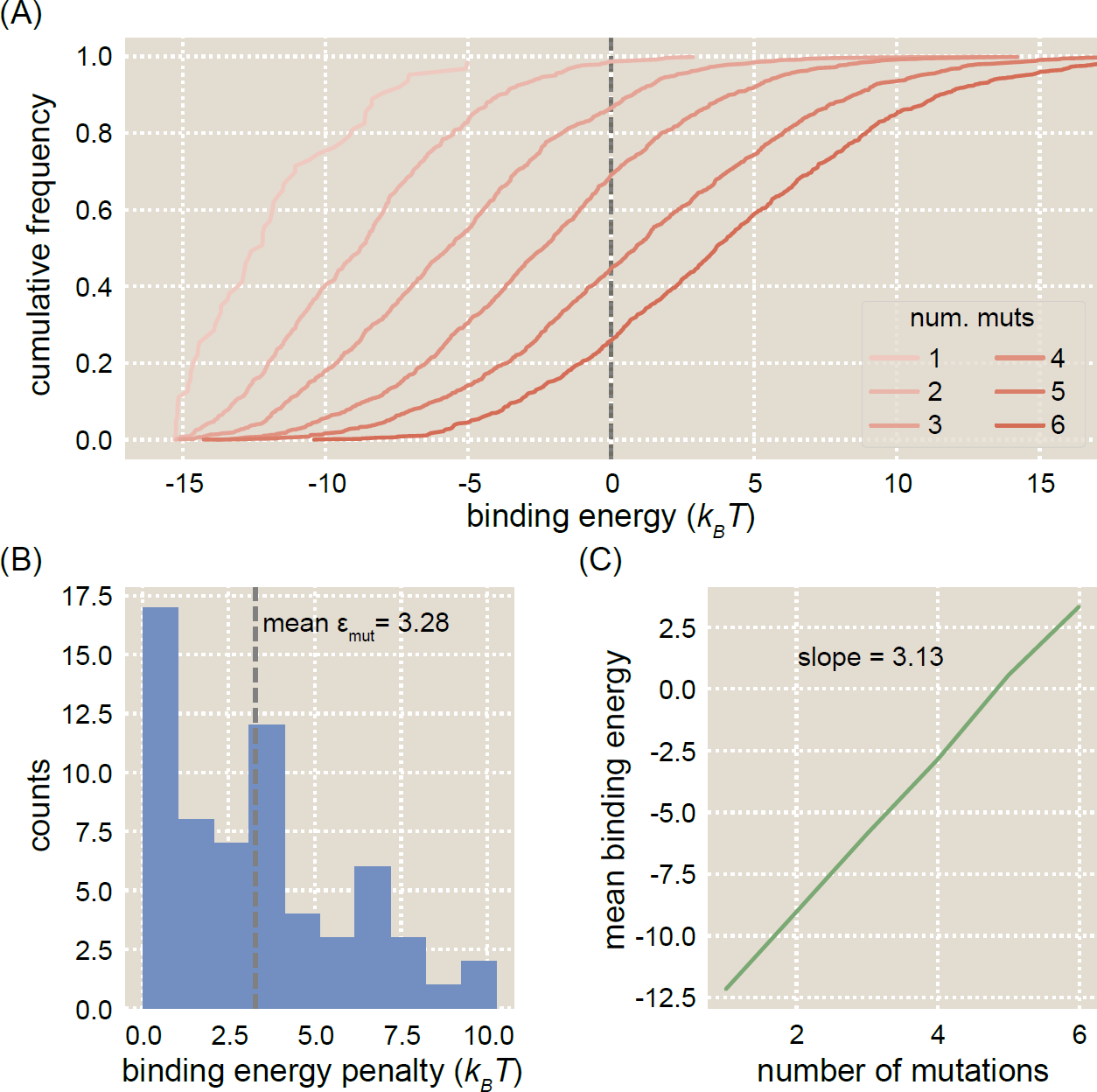
Average effect of a binding site mutation. (A) Cumulative distributions are shown for the predicted binding energies of *lac* operator mutants. The mean predicted binding energy increases substantially with the number of mutations, as does the width of the distribution. The dotted line shows the point at which Δ*ε*_*R*_ = 0 *k*_*B*_*T*, which is the average energy of nonspecific binding. (B) A histogram of binding penalties for single base-pair mutations to the minimum-energy LacI binding sequence shows that the mean binding penalty of a mutation is 3.28 *k*_*B*_*T*. (C) Plotting the mean binding energy of an operator against the number of mutations relative to the minimum-energy sequence shows a linear trend with a slope approximately equal to the average energy penalty per mutation.

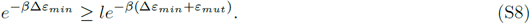

This implies that the minimum average binding energy penalty due to a mutation is given by *ε*_*mut*_ = ln *l*, which for a binding site of 21 bp (the length of a *lac* operator) comes out to *ε*_*mut*_ *≈* 3 *k*_*B*_*T*. This is remarkably close to the mean energy penalty of 3.28 *k*_*B*_*T* calculated for LacI as noted in Figure S2B.

Based on this estimate, one can find a value for *α* by setting the minimum binding energy of an energy matrix to 0, then taking the mean of the nonzero elements of the matrix, *ε*_*mean*_, and finding a scaling factor *α* such that *αε*_*mean*_ = ln *lk*_*B*_*T*.

#### Method comparison

Each of the methods outlined above is capable of producing a value of *α* that can be used to convert an energy matrix into *k*_*B*_*T* units. Each of these methods has its own advantages and disadvantages. Here we will outline these trade-offs and compare the accuracy of the predictions that can be made using each method.

The primary advantage of the Bayesian regression by MCMC method, which is used for the LacI binding energy predictions in the main text, is that it can be implemented using the same Sort-Seq data that was used to obtain the energy matrices. No further data collection is required. However, in order to implement this method one must have a thermodynamic model that predicts gene expression for a given operator binding energy. This is trivial for systems with simple regulatory architectures, as is the case with the simple repression architecture used in this study. However, while models for more complex architectures have been proposed [28], identifying the correct model may not be straightforward and a number of additional experiments may be required in order to validate the proposed model. Additionally, significant computing power is required in order to infer a scaling factor using this method.

The advantages of the least-squares fitting method are that it is conceptually straightforward, it requires little computing power, and it provides a very accurate scaling factor. However, multiple fold-change measurements for different operator mutants are required to perform the regression and calculate the best-fit value of *α*, and any outliers must be identified in order to maximize the accuracy of the fit. Additionally, a thermodynamic model for the system is again required if binding energies are to be measured using fold-change data.

The advantage of the theoretical mutation parameter method is that it is very simple and requires no knowledge of the regulatory architecture of the promoter. All that is needed is an energy matrix for an operator and an estimate of the operator’s length. Indeed, for XylR we lack sufficient information to confidently infer a thermodynamic model of gene expression, so this is the method used to produce energy matrices for this transcription factor (we note that the theoretical mutation parameter method is also used for PurR energy matrices in the main text, though a thermodynamic model is available for PurR [44]). For the *lac* operator it produces a scaling factor that is approximately as accurate as the other inference methods discussed here (see Figure S3). However, this method is based on simplified biophysical arguments, and it is likely that there are a number of regulatory scenarios for which it would not be as successful.

Figure S3 compares predictions made using each method for obtaining a scaling factor. The same matrix was used for each prediction, with O1 as the wild-type sequence and *R* = 130 LacI tetramers in the strain used for Sort-Seq. We find that all methods produce predictions that generally describe the data, but when comparing the mean squared error (MSE) of the predictions, it is clear that some methods perform better than others. Note that elsewhere in the supplement we compare predictions using the Pearson correlation coefficient (*ρ*). We use the MSE here instead because an inaccurate scaling factor will not affect the linear relationship between predictions and measurements, but it will affect the accuracy of the predictions. Thus a set of predictions may have a high *ρ* value corresponding to a strong linear relationship, but still have a high MSE corresponding to inaccurate predictions. The Bayesian parameter estimation (Figure S3A) and least-squares regression (Figure S3B) methods perform nearly identically. However, while the value for *α* that was inferred from the theoretical mutation parameter (Figure S3C) makes predictions that generally describe the data, the MSE values associated with its predictions are notably larger than the other methods, particularly for the 1 bp mutants.

**Figure S3.**
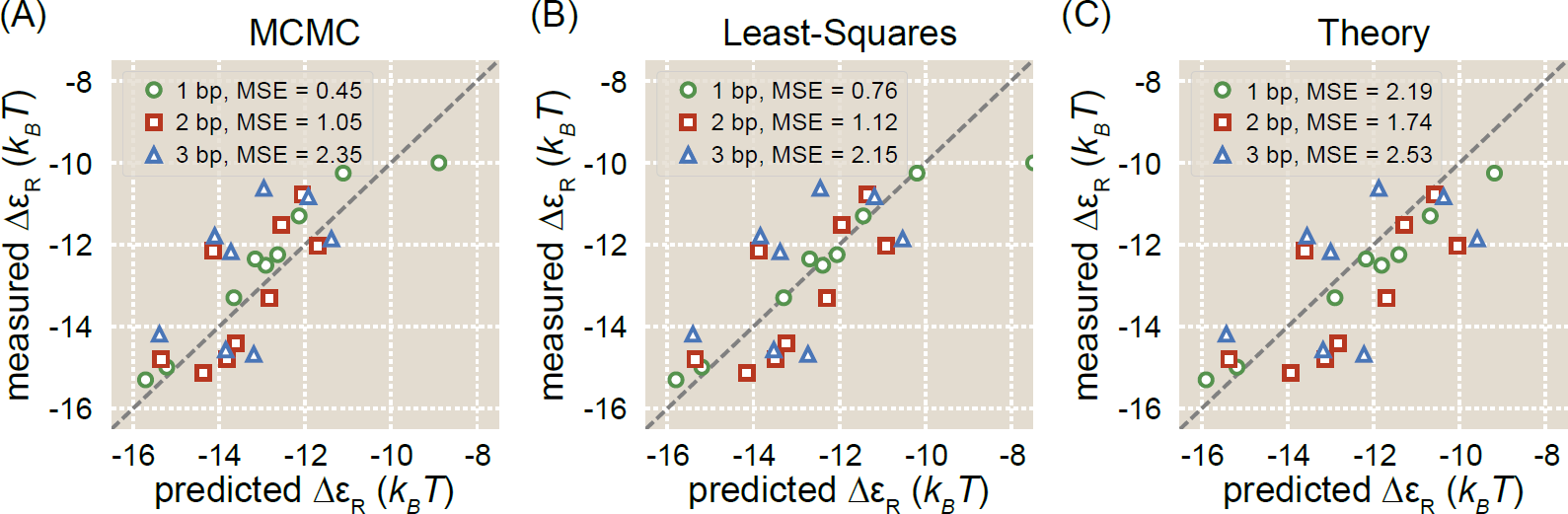
Alternate methods of obtaining energy matrix scaling factor produce similar results. Shown are data for predicted vs. measured binding energies of 1, 2, or 3 bp mutants. The binding energy predictions are made using energy matrices that have been scaled using one of three methods: (A) Bayesian parameter estimation using MCMC, (B) least-squares regression, or (C) inference from a theoretical mutation parameter. All predictions were made using an energy matrix with O1 as the wild-type sequence and *R* = 130 LacI tetramers in the cells used to perform Sort-Seq. The mean squared error (MSE) associated with each set of predictions is noted in the legend.

### D Comparing single-point energy matrix models with higher-order models

A commonly cited problem with the type of energy matrices used in this work is that they do not accurately describe the mechanism of transcription factor binding to DNA. While such energy matrix models assert that each base pair contributes independently to the binding energy, it is known that interactions between two or more base pairs can play an important role in determining binding affinity [33, 66]. In spite of this, energy matrix models that assume independence (which we will refer to here as single-point models) are still commonly used because they often perform nearly as well as higher-order models [22, 67], and they require many fewer parameters than a higher-order model. For example, a single-point model for LacI binding to a 21 bp long operator requires that 84 parameters be inferred, one for each base at each position. By contrast, a two-point model for LacI that accounts for all possible interactions between any two bases in the binding site requires 3660 parameters. Obtaining high-quality estimates for these parameters requires a great deal more data and computing power than inferring parameters for single-point models. Thus it is important to carefully consider whether higher-order models will dramatically improve predictions.

Here we take advantage of our large Sort-Seq data sets to infer two-point binding energy models for LacI binding. As with single-point models, two-point binding energy models are inferred by identifying a set of parameters that maximizes mutual information between sequence and expression bin (see Appendix B for more details). In Figure S4 we compare binding energy measurements to predictions from a single-point model (Fig. S4(A)) and a two-point model (Fig. S4(B)). We also make this comparison for models in which sequences with only one sequencing count are removed from the data set and then all other sequences are weighted equally S4(C-D)). This weighting scheme removes possible sequencing errors from the data set and then gives low-frequency sequences the same influence as high-frequency sequences, compensating for any inequalities that may arise if the library itself has an unequal representation of sequences. The same data set was used to infer each model, namely the data set for the strain with repressor copy number *R* = 130 and an O1 reference sequence. The quality of the predictions for each model is quantified by noting the Pearson’s correlation coefficient *ρ* for each data set. Surprisingly, the unweighted two-point model does not outperform the single-point model. In fact, it performs substantially worse. The weighted two-point model, however, performs better than the weighted single-point model.

**Figure S4.**
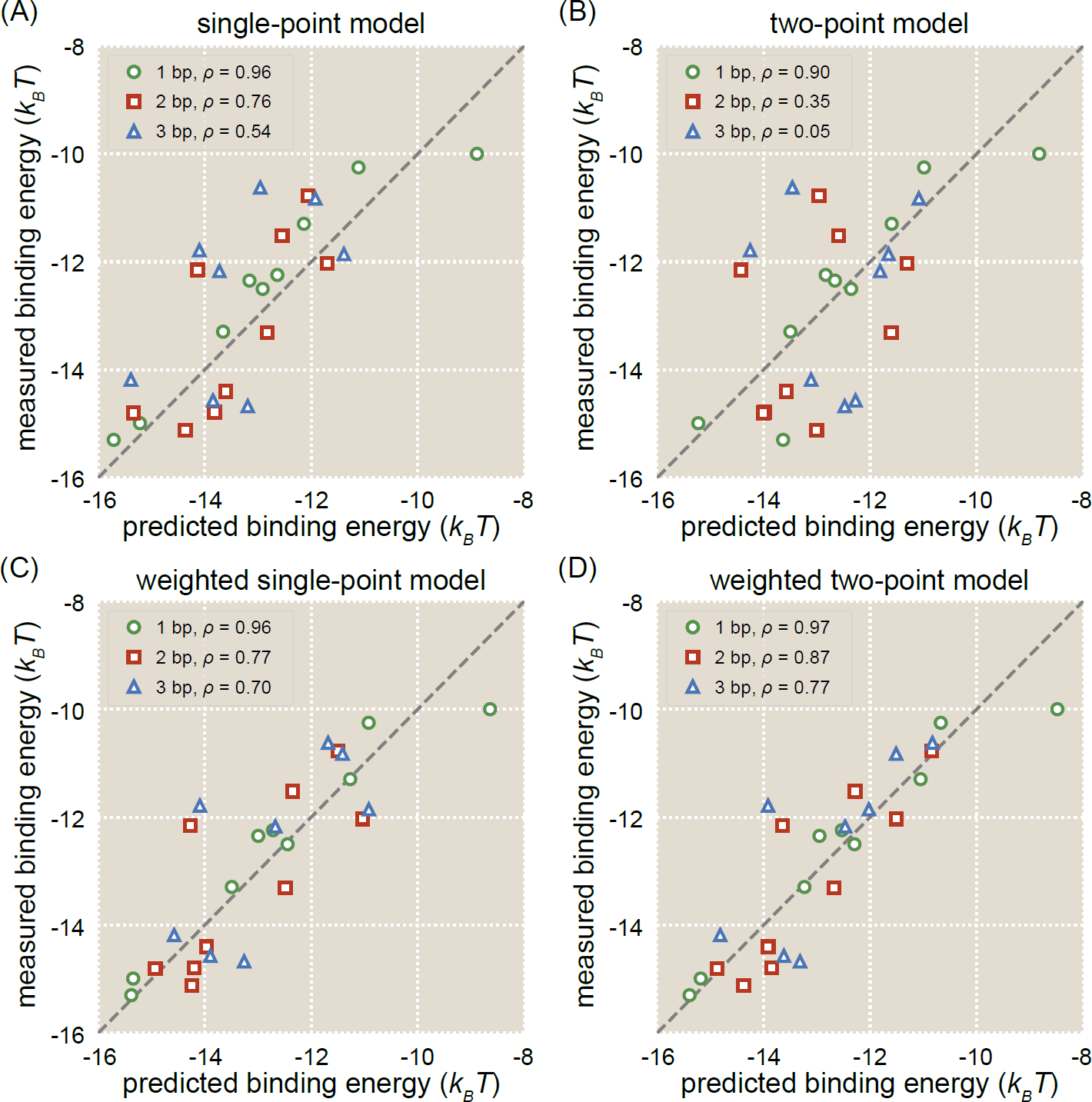
A comparison of single-point models with two-point models. Binding energy measurements are compared against predictions from energy matrix models obtained using a strain where *R* = 130 and O1 is the reference sequence. (A) Predictions are made using a single-point energy matrix in which each sequence position is considered independently. This matrix is used to obtain the predictions discussed in the main text. (B) Predictions are made using an energy matrix model that accounts for all two-point interactions between nucleotides at different sequence positions. The Pearson’s correlation coefficients for the measurements and predictions indicate that this matrix model performs substantially worse than the single-point energy matrix model, particularly for multiple mutations. (C) Predictions are again made using a single-point energy matrix model, though this model has been weighted so that all sequences (aside from single-count sequences, which were dropped) have the same weight. This matrix model has been inferred after removing all single-count sequences from the data set and then weighting all sequences evenly. (D) Predictions are made using a two-point matrix model using the same weighting scheme as in (C). This weighting procedure results in a two-point matrix model that makes improved predictions relative to the weighted single-point energy matrix model.

### E Influence of Regulatory Parameters on Energy Matrix Quality

The level of repression in a repressible system is dependent on a number of factors. In this work we primarily focus on operator binding energy, but other key parameters include operator copy number, repressor copy number, and competition from other binding sites, as discussed in detail in Ref. [68]. Here we consider how two parameters influence energy matrix quality: namely, repressor copy number *R* and the binding energy of the operator reference sequence.

Because our promoter constructs are on plasmids and thus have multiple copies (*N* ≈10), there is some concern that there might not be a sufficient number of repressors in the cell to demonstrate significant changes in expression when the *lac* operator is mutated. The wild-type copy number of LacI tetramers in *E. coli* is *R* = 11, which is comparable to the plasmid copy number used in this study. We increase the LacI copy number by using synthetic RBSs that have been shown to increase gene expression [29]. Additionally, we consider the fact that the binding energy of the reference sequence influences the distribution of binding energies present in the mutant library, and therefore the “ideal” value of *R* may be different for different reference sequences. To explore these factors, we performed Sort-Seq experiments for each combination of *R* (i.e. *R* = 30, 62, 130, or 610) and reference binding energy (i.e. Δ*ε*_R_ = -15.3 *k*_*B*_*T* for O1, Δ*ε*_R_ = -13.9 *k*_*B*_*T* for O2, or Δ*ε*_R_ = -9.7 *k*_*B*_*T* for O3).

#### Comparison of binding energy predictions

Figure S5 shows how predicted and measured binding energy values for single base pair mutants compare for each combination of repressor copy number and reference sequence. We show predictions from energy matrices that have been scaled using the least squares method (see Appendix C), as this is the most accurate method for obtaining a scaling factor. The Pearson’s correlation coefficient (*ρ*) for each set of predictions is shown as a way of quantifying which of these combinations produces the “best” energy matrices, as defined by which matrices give the best agreement between prediction and measurement. We see that the best agreement between prediction and measurement occurs when O1 is the reference sequence. Conversely, predictions from matrices made using O3 as a reference sequence do not predict the measured values at all, as indicated by the especially low *ρ* values. While the choice of repressor copy number does not appear to have a large effect on the quality of matrix predictions, particularly for matrices with O1 as the reference sequence, we do observe that *R* = 610 consistently corresponds with the most accurate predictions. We note that in the main text we make predictions using the energy matrix with the O1 reference sequence and *R* = 130. This is because in the main text we obtain our scaling factors using Bayesian inference by MCMC (see Appendix B), and the most accurate scaling factor inferred by this method was for *R* = 130.

#### Variation in energy matrix replicates

We performed a number of replicates using both O1 and O2 reference sequences to determine the level of variation in sequence logos. As shown in Figure S6, replicates using O1 as a reference sequence produce very consistent sequence logos, while replicates using O2 as a reference sequence produce less consistent sequence logos. This suggests that the strength of the binding site is a significant factor determining the consistency of experiment outcomes.

As another point of comparison, we compared the values making up our energy matrices against one another to assess their consistency. Specifically, we computed the Pearson’s correlation coefficient *ρ* between the lists of values comprising each of our unscaled energy matrices with O1 and O2 reference sequences (see Figure S7). In addition to the matrices analyzed in Figure S5, we performed two additional replicates for each of the energy matrices obtained from strains with *R* = 30 or *R* = 62. This allows us to ascertain whether the matrices themselves are substantially different under different experimental conditions.

We find that all of the matrices with an O1 reference sequence are highly correlated with one another. By contrast, the matrices with an O2 reference sequence are less correlated with one another, even among replicates of the same experimental conditions. The second replicate of the O2 matrix with *R* = 30 is particularly poorly correlated with other matrices. However, the O2 matrices do generally have a higher *ρ* value with one another than with the O1 matrices. An exception to this is the O2 matrices with *R* = 130 and *R* = 610, which appear to be moderately well-correlated with the O1 matrices. These results suggest that the choice of reference sequence used to perform the Sort-Seq experiment is a more important determinant of matrix quality than repressor copy number, though the results may also support the hypothesis that higher repressor copy numbers correspond with improved matrix quality, particularly for weaker reference sequences such as O2.

**Figure S5.**
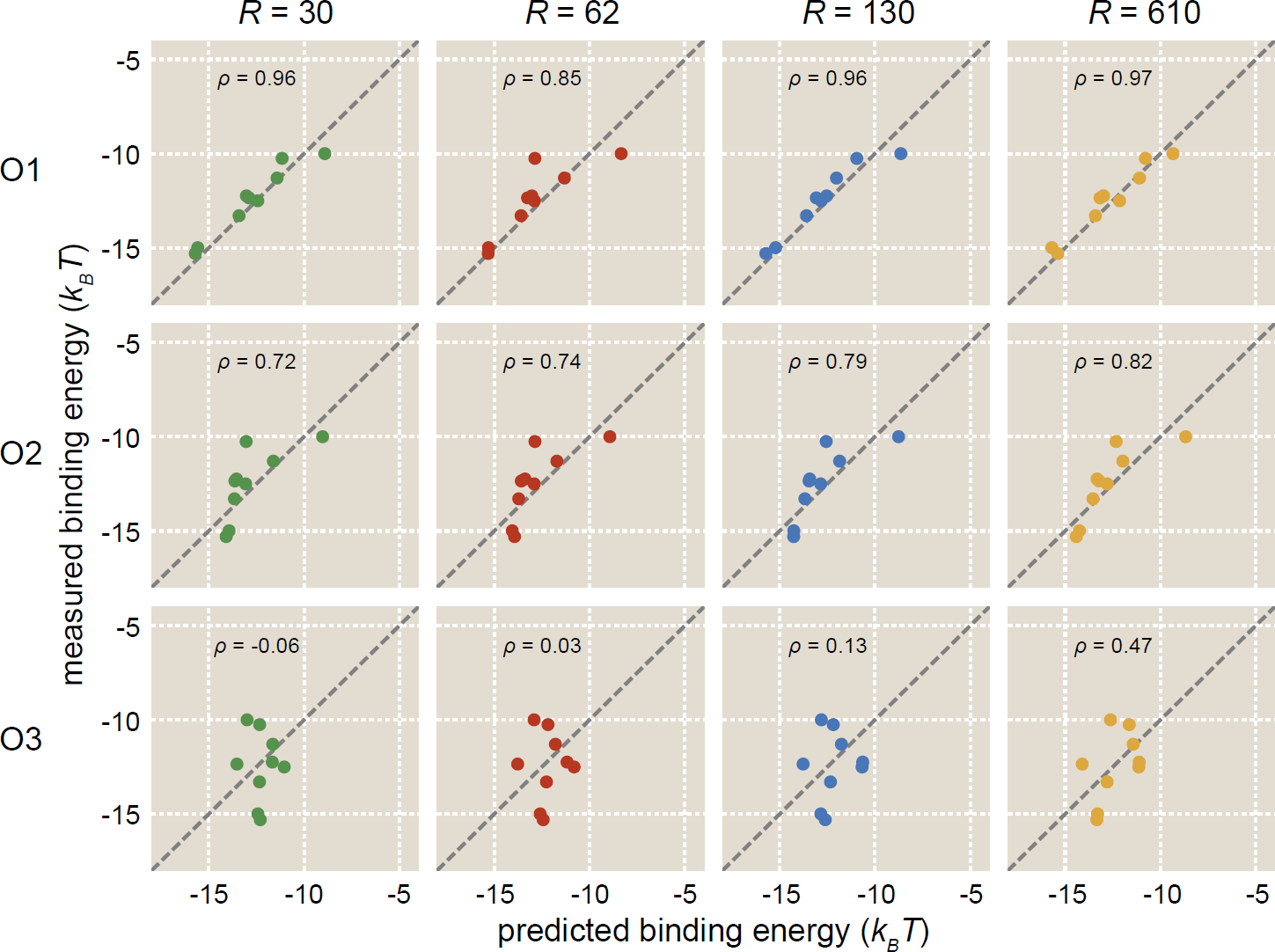
Repressor copy number and reference sequence affect accuracy of energy matrix predictions. Sort-Seq was performed with all combinations of four different repressor copy numbers (*R* = 30, 62, 130, and 610) and three different reference operator sequences (O1, O2, and O3) to produce a total of 12 energy matrices. Predictions from each of these energy matrices are plotted against measured binding energy values for nine single base-pair mutants. The Pearson’s correlation coefficient (*ρ*) is noted for each plot as a measure of prediction accuracy.

**Figure S6.**
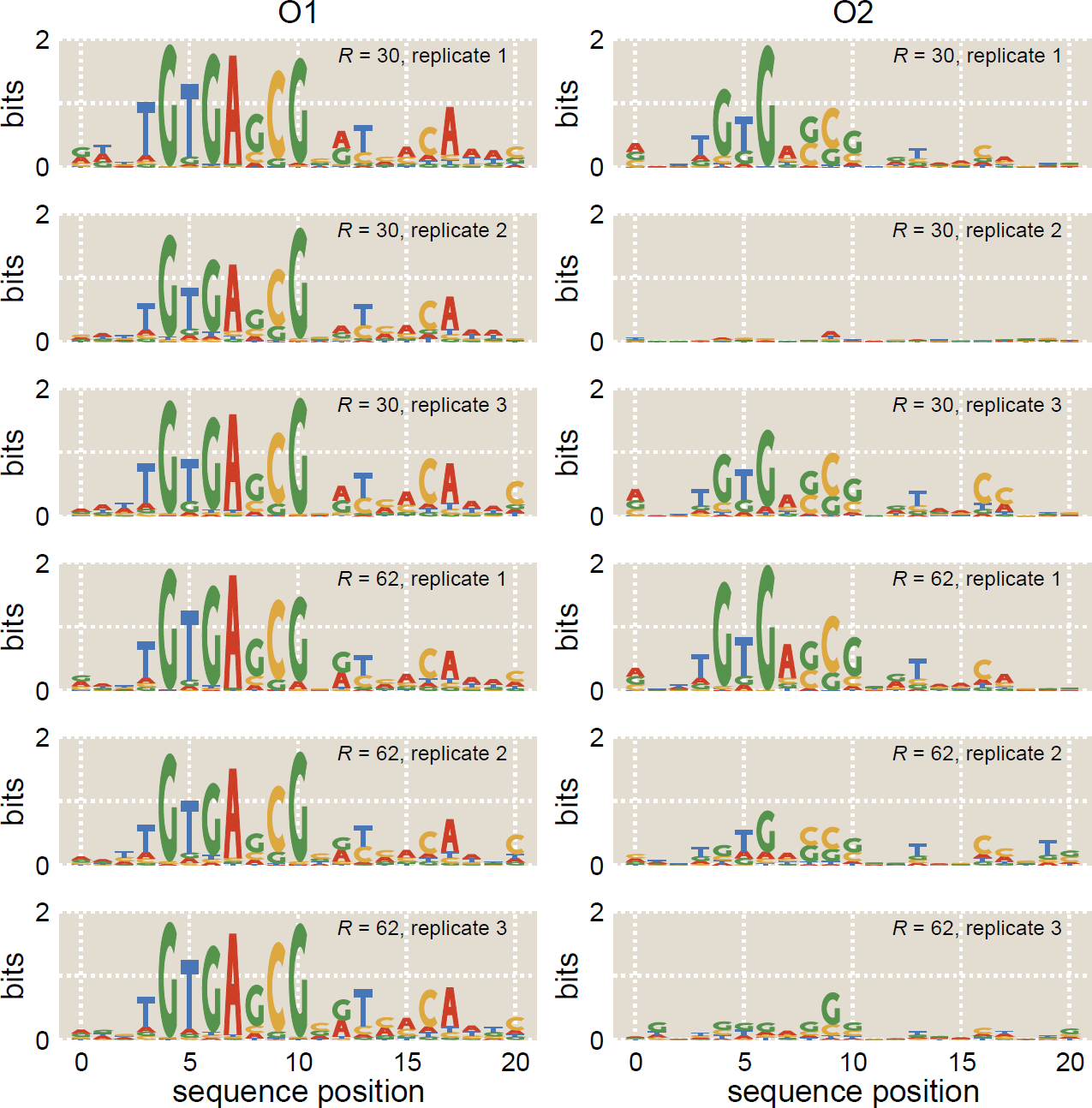
Variation in sequence logo results. Replicates of Sort-Seq experiments were performed using O1 or O2 as a reference sequence. The O1 experiments (left) produced very consistent sequence logos, while the O2 experiments (right) produced sequence logos that varied significantly in quality.

**Figure S7.**
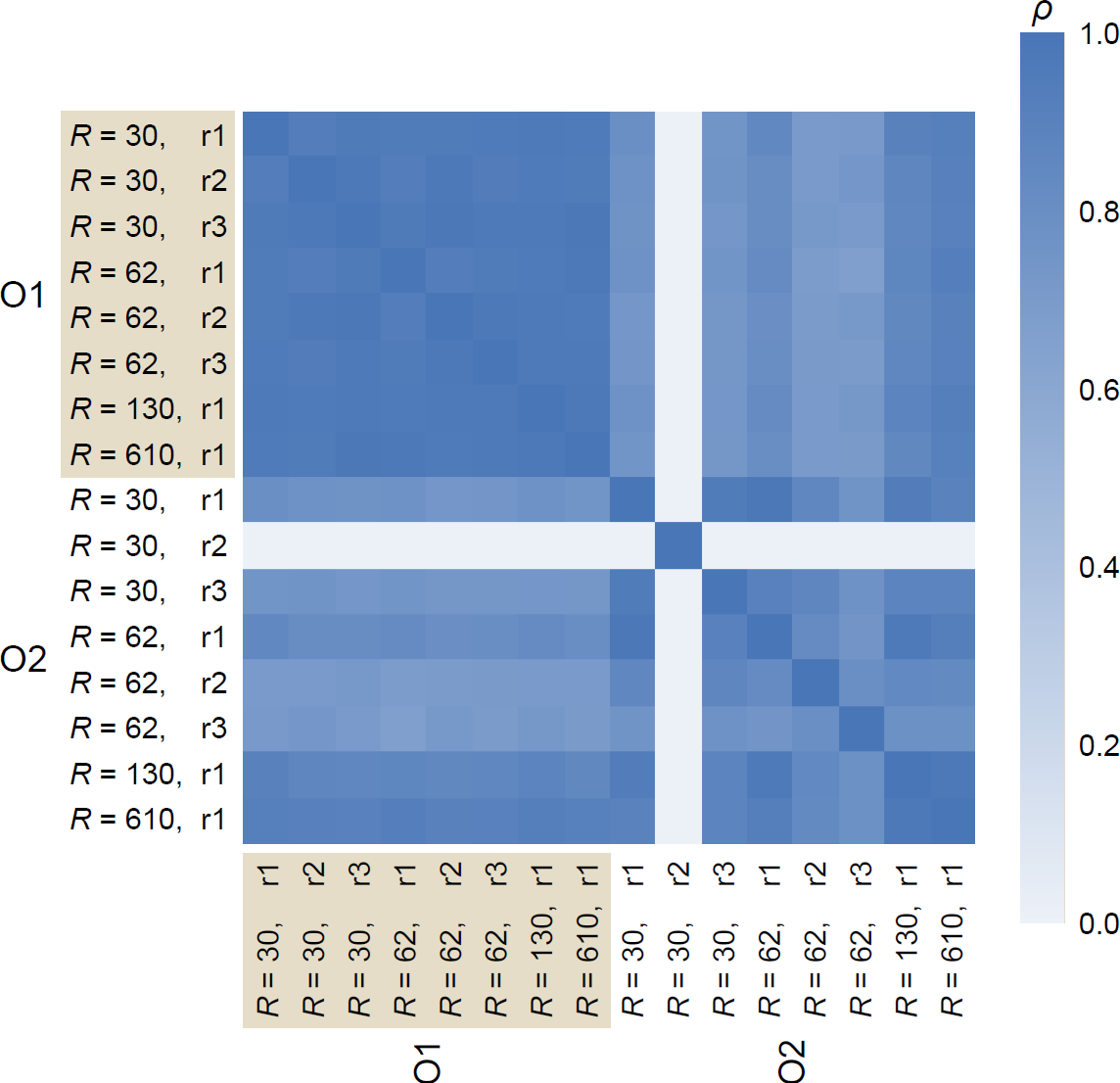
Correlation coefficients between unscaled energy matrices. The Pearson’s correlation coefficient (*ρ*) was calculated for each pair of energy matrices with an O1 or O2 reference sequence. Those experiments conducted using strains with repressor copy number *R* = 30 and *R* = 62 were repeated three times, as denoted by replicate number r1, r2, or r3. We find that all O1 matrices are highly correlated with one another, while O2 matrices are generally less correlated with one another. In general there is low correlation between O1 and O2 matrices, with the exception of O2 matrices with high repressor copy numbers, *R* = 130 and *R* = 610.

### F Comparison of full-promoter and operator-only energy matrix predictions

In the main text, we perform Sort-Seq using libraries in which the entire promoter region was mutated, namely both the RNAP site and the operator. Here we consider whether one can improve energy matrix accuracy by using libraries in which only the operator is mutated.

In order to infer the energy matrix scaling factor *α* from Sort-Seq data alone (see Appendix B), it is necessary to mutate the full promoter, because mutations to both the operator and RNAP binding sites are relevant to the thermodynamic model used to perform the inference. Because of this we use full promoter mutant libraries in the main text. This means that an alternate method is required in order to infer an energy matrix scaling factor for matrices derived from libraries in which only the operator was mutated. Here, we obtain a scaling factor by least-squares regression to a set of measured binding energies for nine 1 bp mutants, as discussed in Appendix C. We then compare measured binding energies against predictions for 1, 2, and 3 bp mutants that were produced using either full-promoter energy matrices or operator-only energy matrices (see Figure S8). We find that operator-only energy matrices produce somewhat more accurate predictions than full-promoter energy matrices. We quantify this by noting the Pearson correlation coefficient (*ρ*) for each set of predictions, which clearly indicates that the O1 operator-only matrix produces the most accurate predictions. This shows us that operator-only energy matrices are a good option when it is feasible to infer the scaling factor from binding energy measurements.

**Figure S8.**
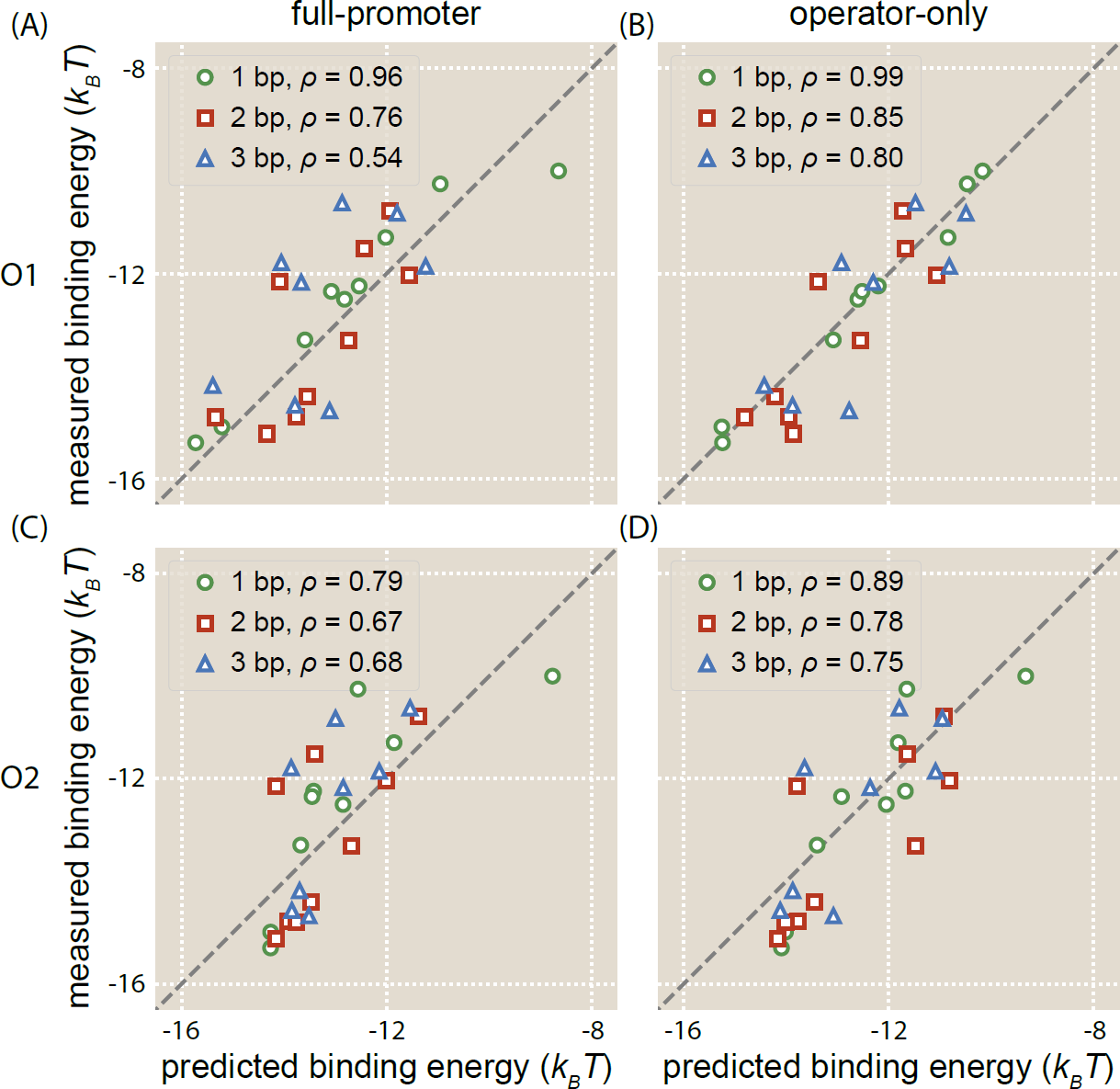
Mutating the operator alone can improve energy matrix accuracy. Binding energy measurements are plotted against energy matrix predictions from full-promoter (A, C) and operator-only (B, D) energy matrices using either O1 (A, B) or O2 (C, D) as a reference sequence. The Pearson correlation coefficient (*ρ*) is noted for each set of predictions. We see that the operator-only energy matrices produce more accurate predictions than the full-promoter energy matrices.

### G Summary of all fold-change data

To measure binding energies for each mutant, fold-change measurements first were obtained by flow cytometry for each mutant in strains with repressor copy numbers *R* = 11 ± 1, 30 ± 10, 62 ± 15, 130 ± 20, 610 ± 80, and 870 ± 170. The data were fit to the fold-change equation

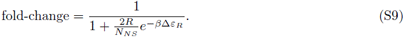

Nonlinear regression was used to obtain the most probable value of Δ*ε*_*R*_ for each mutant. The fold-change data, fitted theory curve, and predicted theory curve are shown here for all 1 bp (Figure S9), 2 bp (Figure S10), and 3 bp mutants (Figure S11).

**Figure S9.**
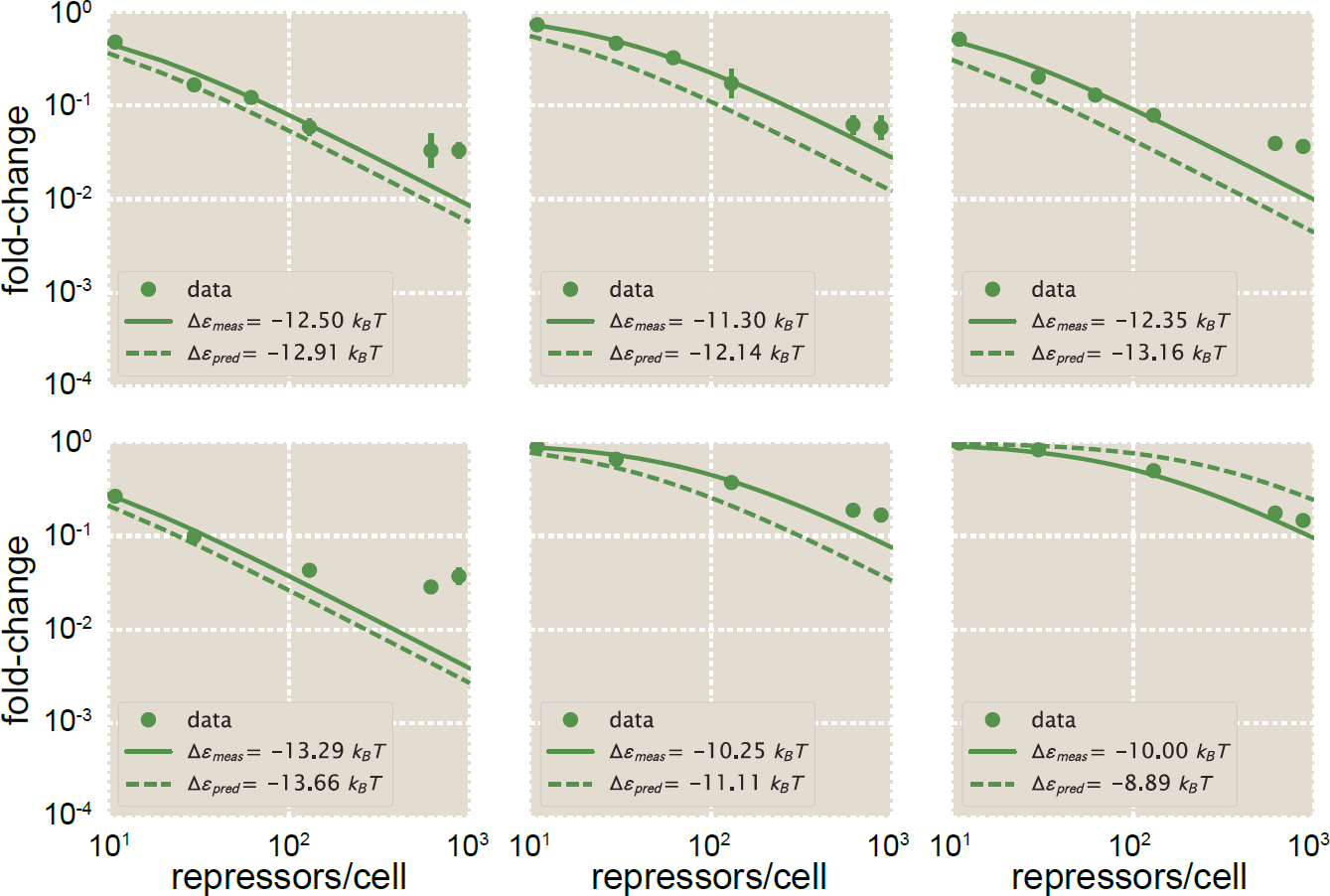
Fold-change measurements for 1 bp mutants. Fold-change measurements are shown for nine 1 bp operator mutants in strains with *R* = 11, 30, 62, 130, 610, or 870. These measurements are overlaid with the measured (fitted) binding energy measurements for each mutant and the predicted measurements as listed in the main text. Note that the bottom three plots do not display data points for *R* = 62, as the data for these strains were outliers.

**Figure S10.**
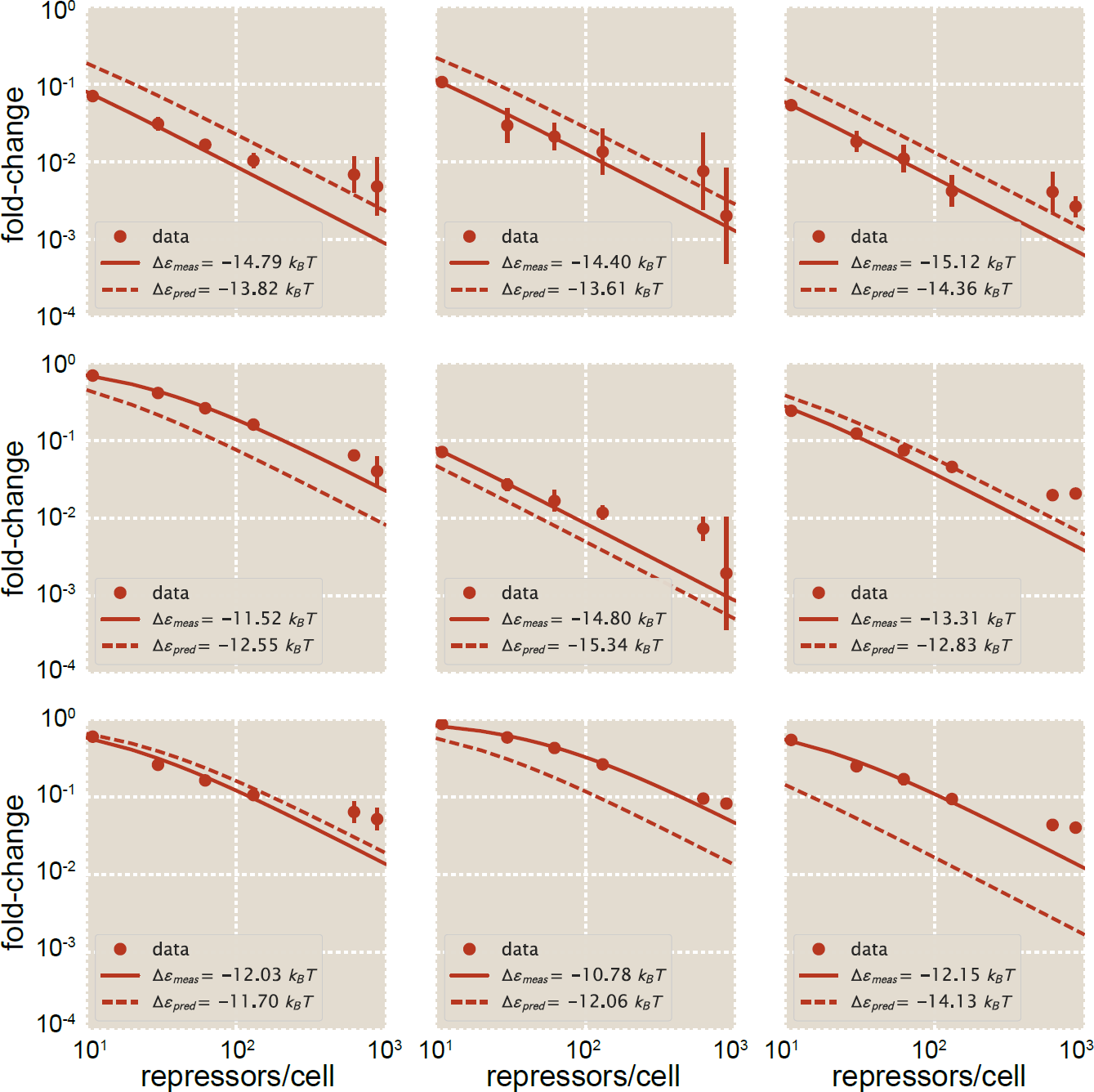
Fold-change measurements for 2 bp mutants. Fold-change measurements are shown for nine 2 bp operator mutants in strains with *R* = 11, 30, 62, 130, 610, or 870. These measurements are overlaid with the measured (fitted) binding energy measurements for each mutant and the predicted measurements as listed in the main text.

**Figure S11.**
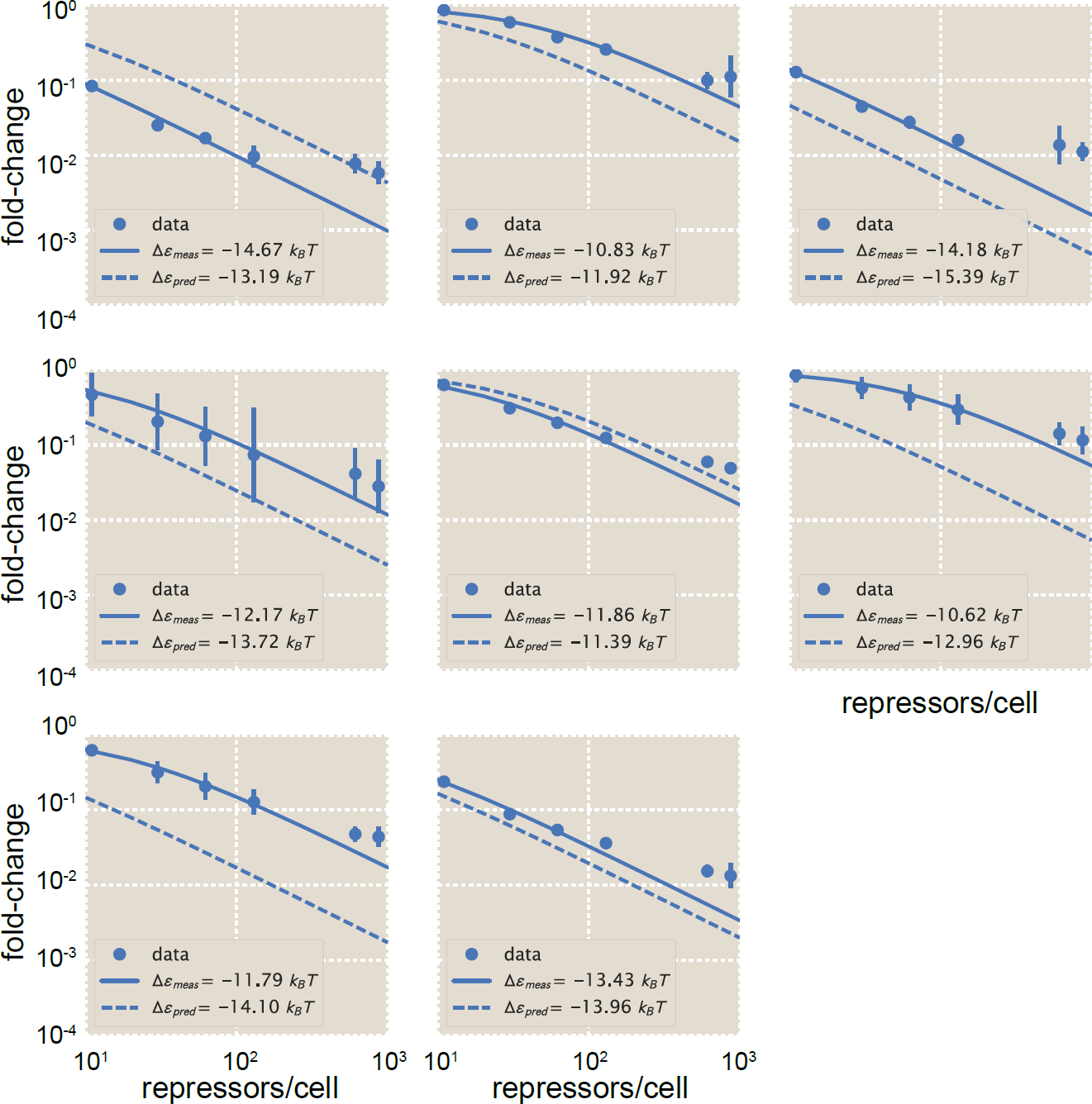
Fold-change measurements for 3 bp mutants. Fold-change measurements are shown for eight 3 bp operator mutants in strains with *R* = 11, 30, 62, 130, 610, or 870. These measurements are overlaid with the measured (fitted) binding energy measurements for each mutant and the predicted measurements as listed in the main text.

### H Expressions for phenotypic parameters of induction responses

As discussed in greater detail in Ref. [41], the thermodynamic model we use to predict induction responses allows us to derive expressions for the phenotypic parameters of the induction response. Here we briefly list the expressions for the phenotypic parameters we address in the present work.

The leakiness of the induction curve is the minimum fold-change observed in the absence of ligand, given by

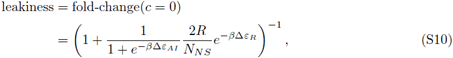

where *c* is the concentration of inducer, *n* is the number of inducer binding sites on the repressor, and Δ*ε*_*AI*_ is the difference in free energy between the repressor’s active and inactive states.

The saturation is the maximum fold change observed in the presence of saturating ligand,

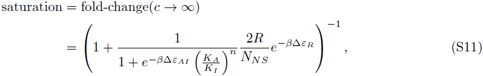

where *K*_*A*_ and *K*_*I*_ are the dissociation constants of the inducer and repressor when the repressor is in its active or inactive state, respectively.

Together, these two properties determine the dynamic range of a system’s response, which is given by the difference

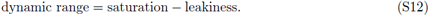

The full expression for dynamic range is then given by

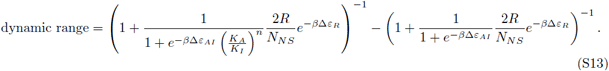

The [*EC*_50_] of the induction response denotes the inducer concentration required to generate a system response halfway between its minimum and maximum value such that

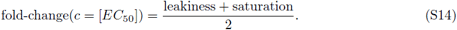

The full expression for the [*EC*_50_] is then given by

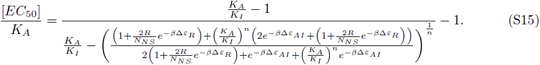

## References

1. Gama-castro S, Salgado H, Santos-zavaleta A, Ledezma-tejeida D, Mu L, Garc JS, et al. RegulonDB version 9.0: high-level integration of gene regulation, coexpression, motif clustering and beyond. Nucleic Acids Research. 2016;44:133–143.

2. Oehler S, Eismann ER, Krämer H, Müller-Hill B. The three operators of the *lac* operon cooperate in repression. The EMBO journal. 1990;9(4):973–979.

3. Gerdes K, Christensen SK, Lobner-Olesen A. Prokaryotic toxin-antitoxin stress response loci. Nature Reviews Microbiology. 2005;3:371–382.

4. Alekshun MN, Levy SB. Regulation of chromosomally mediated multiple antibiotic resistance: The *mar* regulon. Antimicrobial Agents and Chemotherapy. 1997;41(10):2067–2075.

5. Minchin SD, Busby SJW. Analysis of mechanisms of activation and repression at bacterial promoters. Methods. 2009;47(1):6–12.

6. Levo M, Avnit-Sagi T, Lotan-Pompan M, Kalma Y, Weinberger A, Yakhini Z, et al. Systematic investigation of transcription factor activity in the context of chromatin using massively parallel binding and expression assays. Molecular Cell. 2017;65(4):604–617.

7. Melnikov A, Murugan A, Zhang X, Tesileanu T, Wang L, Rogov P, et al. Systematic dissection and optimization of inducible enhancers in human cells using a massively parallel reporter assay. Nature Biotechnology. 2012;30(3):271–277.

8. Levy L, Anavy L, Solomon O, Cohen R, Brunwasser-meirom M, Ohayon S, et al. A synthetic oligo library and sequencing approach reveals an insulation mechanism encoded within bacterial *σ*^54^ promoters. Cell Reports. 2017;21(3):845–858.

9. Berger MF, Philippakis AA, Qureshi AM, He FS, III PWE, Bulyk ML. Compact, universal DNA microarrays to comprehensively determine transcription-factor binding site specificities. Nature Biotechnology. 2006;24(11):1429–1435.

10. Fields DS, He Yy, Al-Uzri AY, Stormo GD. Quantitative specificity of the Mnt repressor. Journal of Molecular Biology. 1997;271:178–194.

11. Jolma A, Yan J, Whitington T, Toivonen J, Nitta KR, Rastas P, et al. DNA-binding specificities of human transcription factors. Cell. 2013;152:327–339.

12. Rastogi C, Rube HT, Kribelbauer JF, Crocker J, Loker RE, Martini GD, et al. Accurate and sensitive quantification of protein-DNA binding affinity. Proceedings of the National Academy of Sciences. 2018;In press.

13. Maerkl SJ, Quake SR. A systems approach to measuring the binding energy landscapes of transcription factors. Science. 2007;315:233–238.

14. Shultzaberger RK, Maerkl SJ, Kirsch JF, Eisen MB. Probing the informational and regulatory plasticity of a transcription factor DNA-binding domain. PLoS Genetics. 2012;8(3).

15. Le DD, Shimko TC, Aditham AK, Keys AM, Orenstein Y, Fordyce P. Comprehensive, high-resolution binding energy landscapes reveal context dependencies of transcription factor binding. Proceedings of the National Academy of Sciences. 2018;In press.

16. Jung C, Hawkins JA, Jones SK, Xiao Y, Rybarski JR, Dillard KE, et al. Massively parallel biophysical analysis of CRISPR-Cas complexes on next generation sequencing chips. Cell. 2017;170:35–47.

17. Nutiu R, Friedman RC, Luo S, Khrebtukova I, Silva D, Li R, et al. Direct measurement of DNA affinity landscapes on a high-throughput sequencing instrument. Nature Biotechnology. 2011;29(7):659–664.

18. Dror I, Golan T, Levy C, Rohs R, Mandel-Gutfreund Y. A widespread role of the motif environment in transcription factor binding across diverse protein families. Genome Research. 2015;25:1268–1280.

19. Levo M, Zalckvar E, Sharon E, Machado ACD, Kalma Y, Lotam-Pompan M, et al. Unraveling determinants of transcription factor binding outside the core binding site. Genome Research. 2015;25:1018–1029.

20. Christensen RG, Gupta A, Zuo Z, Schriefer LA, Wolfe SA, Stormo GD. A modified bacterial one-hybrid system yields improved quantitative models of transcription factor specificity. Nucleic Acids Research. 2011;39(12):1–9.

21. Xu DJ, Noyes MB. Understanding DNA-binding specificity by bacteria hybrid selection. Briefings in Functional Genomics. 2015;14(1):3–16.

22. Weirauch MT, Cote A, Norel R, Annala M, Zhao Y, Riley TR, et al. Evaluation of methods for modeling transcription factor sequence specificity. Nature Biotechnology. 2013;31(2):126–134.

23. Djordjevic M, Sengupta AM, Shraiman BI. A biophysical approach to transcription factor binding site discovery. Genome Research. 2003;13:2381–2390.

24. Garcia HG, Sanchez A, Boedicker JQ, Osborne M, Gelles J, Kondev J, et al. Operator sequence alters gene expression independently of transcription factor occupancy in bacteria. Cell Reports. 2012;2:150–161.

25. Wunderlich Z, Mirny LA. Different gene regulation strategies revealed by analysis of binding motifs. Trends in Genetics. 2009;25(10):434–440.

26. Brewster RC, Jones DL, Phillips R. Tuning promoter strength through RNA polymerase binding site design in *Escherichia coli*. PLoS Computational Biology. 2012;8(12).

27. Kinney JB, Murugan A, Callan CG, Cox EC. Using deep sequencing to characterize the biophysical mechanism of a transcriptional regulatory sequence. Proceedings of the National Academy of Sciences. 2010;107(20):9158–9163.

28. Bintu L, Buchler NE, Garcia HG, Gerland U, Hwa T, Kondev J, et al. Transcriptional regulation by the numbers: models. Current Opinion in Genetics and Development. 2005;15:116–124.

29. Garcia HG, Phillips R. Quantitative dissection of the simple repression input-output function. Proceedings of the National Academy of Sciences. 2011;108(29):12173–12178.

30. Berg OG, Hippel PHV. Selection of DNA binding sites by regulatory proteins: statistical-mechanical theory and application to operators and promoters. Journal of Molecular Biology. 1987;193:723–750.

31. Kalodimos CG, Bonvin AMJJ, Salinas RK, Wechselberger R, Boelens R, Kaptein R. Plasticity in protein-DNA recognition: *lac* repressor interacts with its natural operator O1 through alternative conformations of its DNA-binding domain. The EMBO Journal. 2002;21(12):2866–2876.

32. Zuo Z, Stormo GD. High-resolution specificity from DNA sequencing highlights alternative modes of *lac* repressor binding. Genetics. 2014;198:1329–1343.

33. Siebert M, Söding J. Bayesian Markov models consistently outperform PWMs at predicting motifs in nucleotide sequences. Nucleic Acids Research. 2016;44(13):6055–6069.

34. Rogers JK, Guzman CD, Taylor ND, Raman S, Anderson K, Church GM. Synthetic biosensors for precise gene control and real-time monitoring of metabolites. Nucleic Acids Research. 2015;43(15):7648–7660.

35. Rohlhill J, Sandoval NR, Papoutsakis ET. Sort-seq approach to engineering a formaldehyde-inducible promoter for dynamically regulated *Escherichia coli* growth on methanol. ACS Synthetic Biology. 2017;6(8):1584–1595.

36. Moon TS, Lou C, Tamsir A, Stanton BC, Voigt CA. Genetic programs constructed from layered logic gates in single cells. Nature. 2012;491:249–253.

37. Collins JJ, Gardner TS, Cantor CR. Construction of a genetic toggle switch in *Escherichia coli*. Nature. 2000;403:339–342.

38. Ozbudak EM, Thattai M, Kurtser I, Grossman AD, Oudenaarden AV. Regulation of noise in the expression of a single gene. Nature Genetics. 2002;31:69–73.

39. Rosenfeld N, Young JW, Alon U, Swain PS, Elowitz MB. Gene regulation at the single-cell level. Science. 2005;307:1962–1965.

40. Purnick PEM, Weiss R. The second wave of synthetic biology: from modules to systems. Nature Reviews Molecular Cell Biology. 2009;10:410–422.

41. Razo-Mejia M, Barnes SL, Belliveau NM, Chure G, Einav T, Lewis M, et al. Tuning transcriptional regulation through signaling: A predictive theory of allosteric regulation. Cell Systems. 2018;In press.

42. Milk L, Daber R, Lewis M. Functional rules for *lac* repressor-operator associations and implications for protein-DNA interactions. Protein Science. 2010;19:1162–1172.

43. Daber R, Sochor MA, Lewis M. Thermodynamic analysis of mutant *lac* repressors. Journal of Molecular Biology. 2011;409:76–87.

44. Belliveau NM, Barnes SL, Ireland WT, Jones DL, Sweredoski M, Moradian A, et al. A systematic approach for dissecting the molecular mechanisms of transcriptional regulation in bacteria. Proceedings of the National Academy of Sciences. 2018;In press.

45. Rohs R, West SM, Sosinsky A, Liu P, Mann RS, Honig B. The role of DNA shape in protein–DNA recognition. Nature. 2009;461:1248–1253.

46. Rohs R, Jin X, West SM, Joshi R, Honig B, Mann RS. Origins of specificity in protein-DNA recognition. Annual Reviews in Biochemistry. 2010;79:233–269.

47. Slattery M, Riley T, Liu P, Abe N, Gomez-alcala P, Dror I, et al. Cofactor binding evokes latent differences in DNA binding specificity between Hox proteins. Cell. 2011;147:1270–1282.

48. Rydenfelt M, Garcia HG, Cox RS, Phillips R. The influence of promoter architectures and regulatory motifs on gene expression in *Escherichia coli*. PLoS ONE. 2014;9(12):1–31.

49. Brophy JAN, Voigt CA. Principles of genetic circuit design. Nature Methods. 2014;11(5):508–520.

50. Khalil AS, Collins JJ. Synthetic biology: applications come of age. Nature Reviews Genetics. 2010;11:367–379.

51. Bréchemier-Baey D, Domínguez-RaMírez L, Plumbridge J. The linker sequence, joining the DNA-binding domain of the homologous transcription factors, Mlc and NagC, to the rest of the protein, determines the specificity of their DNA target recognition in *Escherichia coli*. Molecular Microbiology. 2012;85(5):1007–1019.

52. Camas FM, Alm EJ, Poyatos JF. Local gene regulation details a recognition code within the LacI transcriptional factor family. PLoS Computational Biology. 2010;6(11).

53. Urtecho G, Tripp AD, Insigne K, Kim H, Kosuri S. Systematic dissection of sequence elements controlling *σ*^70^ promoters using a genomically-encoded multiplexed reporter assay in *E. coli*. Biochemistry. 2018;In press.

54. Patwardhan RP, Lee C, Litvin O, Young DL, Pe’Er D, Shendure J. High-resolution analysis of DNA regulatory elements by synthetic saturation mutagenesis. Nature Biotechnology. 2009;27(12):1173–1175.

55. Sharon E, Kalma Y, Sharp A, Raveh-Sadka T, Levo M, Zeevi D, et al. Inferring gene regulatory logic from high-throughput measurements of thousands of systematically designed promoters. Nature Biotechnology. 2012;30(6):521–530.

56. Smith RP, Taher L, Patwardhan RP, Kim MJ, Inoue F, Shendure J, et al. Massively parallel decoding of mammalian regulatory sequences supports a flexible organizational model. Nature Genetics. 2013;45(9):1021–1028.

57. Lutz R, Bujard H. Independent and tight regulation of transcriptional units in *Escherichia coli* via the LacR/O, the TetR/O and AraC/I1-I2 regulatory elements. Nucleic Acids Research. 1997;25(6):1203–1210.

58. Salis HM, Mirsky EA, Voigt CA. Automated design of synthetic ribosome binding sites to control protein expression. Nature Biotechnology. 2009;27(10):946–950.

59. Datta S, Costantino N, Court DL. A set of recombineering plasmids for gram-negative bacteria. Gene. 2006;379:109–115.

60. Kinney J, Atwal GS. Parametric inference in the large data limit using maximally informative models. Neural Computation. 2014;26(4):637–653.

61. Kinney JB, Tkačik G, Callan CG. Precise physical models of protein-DNA interaction from high-throughput data. Proceedings of the National Academy of Sciences. 2007;104(2):501–506.

62. Atwal GS, Kinney JB. Learning quantitative sequence–function relationships from massively parallel experiments. Journal of Statistical Physics. 2016;162(5):1203–1243.

63. Ireland WT, Kinney JB. MPAthic: quantitative modeling of sequence-function relationships for massively parallel assays. bioRxiv. 2016;.

64. Earl DJ, Deem MW. Parallel tempering: Theory, applications, and new perspectives. Physical Chemistry Chemical Physics. 2005;7:3910–3916.

65. Lässig M. From biophysics to evolutionary genetics: statistical aspects of gene regulation. BMC Bioinformatics. 2007;8(Suppl 6):S7.

66. Benos PV, Bulyk ML, Stormo GD. Additivity in protein-DNA interactions: how good an approximation is it? Nucleic Acids Research. 2002;30(20):4442–4451.

67. Zhao Y, Stormo GD. Quantitative analysis demonstrates most transcription factors require only simple models of specificity. Nature Biotechnology. 2011;29(6):480–483.

68. Weinert FM, Brewster RC, Rydenfelt M, Phillips R, Kegel WK. Scaling of gene expression with transcription-factor fugacity. Physical Review Letters. 2014;113.

